# Epigenetic Gatekeeping of Wnt-Responsive Enhancers Restricts Intestinal Stem Cell Transformation

**DOI:** 10.64898/2026.01.31.702974

**Authors:** Alireza Lorzadeh, Sweta Sharma, George Ye, Sabrina Rabaya, Yun Jee Kang, Ramesh A. Shivdasani, Unmesh Jadhav

## Abstract

Widespread cell plasticity recognized in fetal intestinal epithelium is preserved in limited fashion in Wnt-responsive adult stem cells and contributes to tumour initiation, progression, and relapse. It is unclear which epigenetic features maintain stem-cell properties, restrict adult expression of fetal genes, and are attenuated in tumours, allowing non-stem cells to replenish targeted tumour stem cells. Here we show that reversible stemness in normal adult intestinal crypt cells hinges on a dynamic balance between activating H3K27ac and repressive H3K27me3 marks. Cells that leave the Wnt-rich stem-cell niche normally acquire H3K27me3 at thousands of stemness-associated enhancers. Constitutive tumourigenic Wnt activity transforms *Apc^‒/‒^* intestinal stem cells by gradual erosion of H3K27me3 at select enhancers and extends stem-like properties beyond usual anatomic confines; continued depletion of H3K27me3 reactivates enhancers that control growth and expression of a wider swath of fetal genes than appreciated previously. Subsequent focal DNA demethylation at expanded superenhancer domains is associated with tumour growth. Human colorectal cancers also carry evidence of this epigenetic rewiring. Accelerated H3K27me3 loss in mice hastens, and its preservation delays, activation of stemness-related enhancers, superenhancers, and tumour progression. During transformation, H3K27me3 loss at enhancers erases a crucial distinction between stem and non-stem populations, endowing the latter with stemness and providing an explanation for tumour resistance to cancer stem cell targeting. Thus, H3K27me3 at Wnt-responsive enhancers is an intrinsic barrier to intestinal tumourigenesis and aberrant reactivation of hundreds of fetal genes.

---

Adult intestinal stem cells (ISCs) generate millions of absorptive and secretory progeny daily^1^. Stemness, the combined capacity for self-renewal and multilineage potential, is confined to Lgr5^+^ cells at the base of epithelial crypts^1^ and maintained in part by Wnt–Rspondin signaling^2^. During fetal life, Lgr5^+^ and Lgr5^‒^ epithelial cells can interconvert and generate adult stem cells^3^. In adults, *Lgr5* expression peaks at crypt bottoms and declines upward (Fig. 1a), reflecting a high-Wnt niche that regulates stem-like properties through epithelial cell movement.^1, 4, 5 1, 6^ (Fig. 1a). Stochastic return of some cells to the crypt bottom reinstates stemness^7^, emphasizing its dynamic quality and implying precise regulation of the balance between activity and silencing of stemness-related transcriptional programs.

**Figure 1.**
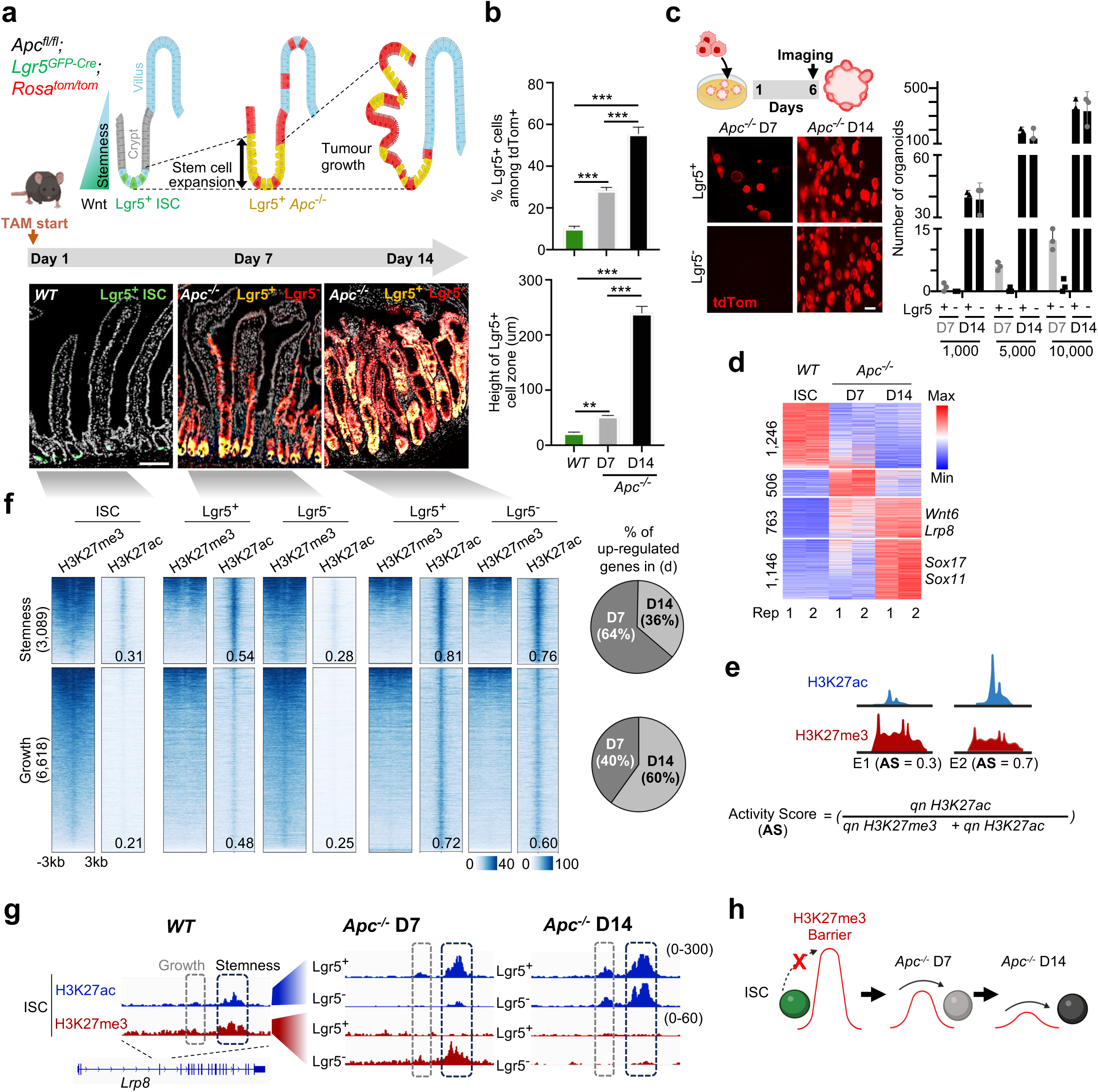
H3K27me3-based barrier confines stemness to intestinal stem cells. **a.** Early events following Apc deletion: Lgr5^+^ ISCs expand by day 7 (D7), increasing stem cell zone height, and progressively spreading throughout the growing adenoma day 14 (D14). Representative immunofluorescence images show expansion of transformed (*Apc^‒/‒^*) Lgr5^+^ stem cells (yellow) in crypts versus wild-type (green), and emergence of their transformed Lgr5⁻ progeny (tdTom^+^, red). Scale bar, 100 μm. **b.** Quantification of increased proportion of Lgr5^+^ cells among tdTom⁺ transformed cells **(top)** and expanded vertical distribution of Lgr5^+^ cells **(bottom)** at D7 and D14 (mean ± s.e.m.; n > 3 mice; \*\**p* < 0.01; \*\*\**p* < 0.001). **c.** Schematic of organoid formation assay and representative images of organoids formed by FACS-isolated Lgr5^+^ and Lgr5^‒^ transformed cells at D7 and D14 (10,000 cells per dome). Scale bar = 50um. Numbers of organoids formed upon seeding 1,000, 5,000, and 10,000 cells. Lgr5^‒^ cells gain organoid forming capacity by D14, approximating the capacity of transformed Lgr5^+^ cells (mean ± s.e.m.; n = 3; ***p < 0.001). **d**. Heatmap showing expression (log2RPKM+1) of differential expressed genes (expression values from 2 replicates) across ISCs and *Apc^‒/‒^* Lgr5^+^ cells at D7 and D14. **e**. Schematic of enhancer activity score calculation. **f.** Heatmap of chromatin remodeling at stemness and growth enhancers across ISCs, villus cells, and *Apc^‒/‒^* Lgr5^+^/Lgr5^‒^ populations at D7 and D14; numbers indicate mean AS values for enhancers in the cluster. Pie charts (right panel) show percentage of up-regulated genes linked to these groups of enhancers at D7 and D14. **g.** Genome browser tracks showing progressive gain of H3K27ac and loss of H3K27me3 at representative stemness (black dashed box) and growth (gray dashed box) enhancers at the *Lrp8* locus. **h.** Model: H3K27me3 restricts stemness-linked enhancer activation to ISCs, while stepwise loss of this barrier during Apc loss-driven transformation enables propagation of stemness and proliferative potential, promoting tumour initiation.

Loss of this precise spatial, niche-level control allows transformed stem cells to expand in intestinal tumours^8–12^. Although niche signals that govern stemness are well characterized^1^, cell-intrinsic mechanisms that safeguard ISC identity are poorly defined and epigenetic mechanisms that control ISC heterogeneity or their homeostatic and transformed states remain unclear. Epigenetic “gatekeeping” is presumed to restrict stemness^13^, but key questions are unresolved: what molecular barriers restrict stemness and when are they established? Does fetal cell plasticity reflect absence of putative barriers? Does their failure extend stem potential to other tumour cells? Lgr5^+^ cells serve as a major source of cancer stem cells (CSCs), sustaining tumour growth.^2^ In tumours, stemness extends beyond the crypt base and Lgr5^‒^ cells acquire stem-like properties^2, 14^, suggesting that transformation dismantles cell-intrinsic barriers. Extended stemness has important consequences in that metastatic Lgr5^‒^ cells engender stem cells at secondary sites^12^ and Lgr5^‒^ cells readily replenish ablated CSCs, driving tumour relapse.^15^ Understanding how tumours breach stem cell-intrinsic molecular barriers is essential for developing strategies that can eradicate CSCs and prevent other cells from acquiring stem-like properties.

Loss of *Apc* function, a regulator of Wnt signaling, occurs in ∼80% of sporadic colorectal cancers (CRCs) and drives constitutive Wnt activity^16, 17^. Because restored *Apc* function ablates even tumours with additional mutations^18^, epigenetic reprogramming of intrinsic stemness must underlie *Apc^‒/‒^* cell transformation. Here we show that H3K27me3 marking at enhancers restricts stemness in adult intestines and steadily erodes during tumourigenesis. The resulting chromatin remodeling and enhancer activation drive stemness in Lgr5^‒^ cells and the spread of fetal programs that accompany tumour initiation and progression. Thus, placement of the repressive H3K27me3 mark at enhancers by the Polycomb complex is a central epigenetic mechanism that restricts expansion, maintenance, and persistence of cancer stemness.

## RESULTS

### Cell growth and expanded stemness occur sequentially in intestinal tumourigenesis

Expanded stemness and uncontrolled growth are hallmarks of tumourigenesis, but how these properties emerge over time and the molecular mechanisms that coordinate them remain poorly understood. To address this gap, we induced adenomas in mouse intestines and examined changes over time in transformed stem and non-stem cell populations. In mammalian intestinal crypts, only a few Lgr5^+^ cells maintain stemness, which is rapidly suppressed as those cells exit the Wnt-rich niche^1, 6^ (Fig. 1a). In *Lgr5^EGFP-CreER-T2^*;*Apc^Fl/Fl^*;*Rosa26^LsL-tdTomato^* mice, injection of tamoxifen on 2 consecutive days induced *Apc* deletion in Lgr5^+^ stem cells (also labeled by EGFP), resulting in intestinal adenomas and tdTomato (tdTom) labeling of mutant cells and their progeny (Fig. 1a, Extended Data Fig. 1a-d). Although mRNA-seq analysis of purified Lgr5^+^ cells (EGFP^+^ tdTom^+^) and their Lgr5^‒^ progeny (EGFP^-^ tdTom^+^) confirmed excision of *Apc* exon 14 by day (D) 7 post-induction (Extended Data Fig. 1c) and the Lgr5^+^ cell zone was expanded, crypt–villus architecture was preserved at this time (Fig. 1a–b, Extended Data Fig. 1e,f). MKi67, a marker of cell proliferation, and nuclear β-catenin, indicating active Wnt signaling, were restricted to Lgr5^+^ cells (Extended Data Fig. 1g). Only by D14 was crypt–villus morphology disrupted, Lgr5^+^ cells extended beyond crypt-like structures, differentiated enterocytes and goblet cells were depleted, and MKi67 and nuclear β-catenin had spread to Lgr5^‒^ progeny (Fig. 1a, Extended Data Fig. 1g-i). To verify the temporal spread of stemness, we assessed self-renewal and proliferative capacity in intestinal organoid cultures (Fig. 1c). We plated FACS-purified Lgr5^+^ and Lgr5^‒^ cells (Extended Data Fig. 1j,k) from D7 and D14 intestines at 1,000, 5,000, or 10,000 cells per dome in medium lacking Rspondin, which permits *Apc^‒/‒^* cell growth^19^. Whereas Lgr5^+^ cells from D7 or D14 readily formed large spherical organoids, Lgr5^‒^ cells from D7 failed to do so, indicating minimal stem potential. By D14, however, Lgr5^‒^ cells formed organoids at densities comparable to Lgr5^+^ cells (Fig. 1c, Extended Data Fig. 1k), revealing that constitutive Wnt activity first unlocks stemness-related properties in a limited cell population and later extends those properties into a growing number of daughter cells, obscuring their functional distinction from the stem-cell compartment. Compared to non-transformed (wild-type, *WT)* Lgr5^+^ ISCs, *Apc^‒/‒^* ISCs upregulated stemness genes, e.g., *Wnt6* and *Lrp8*, by D7; other genes associated with growth, suppressed differentiation, and tissue remodeling, e.g., *Sox11*, *Sox17*, and *Foxc1*, were not upregulated on D7 but active by D14 (Fig. 1d and Extended Data Fig. 1l). Thus, like expansion of stemness, transcriptional dysregulation in stem cells also progresses over a finite period. However, there is no molecular explanation for this stepwise deregulation of genes and cell properties.

### H3K27me3 gatekeeping of intestinal stem cell properties

Because enhancers control cell state–specific genes, we examined how opposing histone modifications associated with cis-element activity (H3K27ac) or inactivity (H3K27me3) influence ISC transformation. We used native-ChIP-seq^20^ to define the distributions of these histone marks in FACS-purified ISCs and in Lgr5^+^ and Lgr5^‒^ cells from D7 and D14 epithelia (Extended Data Fig. 1j). H3K27ac is well-characterized at promoters and enhancers ^21^, but H3K27me3 has been studied principally at promoters and in heterochromatin ^21^. We identified 25,987 genomic sites with either H3K27ac or H3K27me3 marking in *WT* and *Apc^‒/‒^* stem cells, and 13,831 *WT* enhancers where *both marks* were present (Extended Data Fig. 2a-b). To quantify their relative levels and dynamics during transformation, we developed a comparative enhancer “activity score” that considers both signals, appropriately normalized (Fig. 1e, Extended Data Fig. 2a-c; Methods). Scores in the schema range from 0 (H3K27me3 only) to 1 (H3K27ac only) and enhancers with both marks score between 0.3 and 0.7 (Extended Data Fig. 2a, d, e). This approach identified 3,089 ‘*stemness’* enhancers (balanced in *WT*, 0.3 < score < 0.7), which lost H3K27me3 and became notably acetylated (mean score 0.54) in *Apc^‒/‒^* stem cells by D7 but stayed unacetylated (mean score 0.28) in Lgr5^‒^ progeny; by D14, their mean H3K27ac score in Lgr5^‒^ cells also increased to 0.76 (Fig. 1f,g). Additionally, 6,618 ‘*neo’* enhancers lacked H3K27ac in *WT* (score <0.3) and showed modest (mean score 0.48) to no acetylation (mean score 0.25) at D7 in stem and non-stem cells, respectively, but achieved high scores of 0.72 and 0.6 in both compartments by D14 (Δ_score_ >0.3; *p* <0.0001; Fig. 1f,g, Extended Data Fig. 2e,f). Genes near *stemness* enhancers (±25 kb from transcription start site, TSS; e.g., *Lrp8*, *Wnt10a*, *Wnt6, Foxa2*) were enriched for Wnt and cell fate pathways, while those near *neo* enhancers (e.g., *Sox17*, *Sox11*, *Foxc1* – Fig. 1f, Extended Data Fig. 2g-i) were predominantly linked to cell proliferation, signaling, and migration, leading us to consider this class as ‘*growth’* enhancers. The stepwise activation of stemness and growth enhancers are also mirrored in the transcription of their associated genes*, as Apc^‒/‒^* stem cells expressed genes linked to *stemness* enhancers by D7 and those linked to *growth* enhancers only later, by D14 (Fig. 1f, Extended Data Fig. 2j). Although gene activation was initially restricted to Lgr5^+^ cells, the mRNA profiles of Lgr5^+^ and Lgr5^‒^ cells converged by D14 (Extended Data Fig. 2k), coincident with H3K27me3 loss at thousands of enhancers and extension of functional stemness over the same interval (Fig. 1f). Thus, a stemness-related transcriptional program associated with H3K27me3 loss expresses early in ISC transformation and continued erosion of that mark is associated with subsequent gene activity related to cell growth.

Because a given H3K27 residue can be acetylated or methylated, but not both, balanced signals (0.3 < activity score < 0.7, Fig. 1f) at *WT stemness* enhancers likely reflect ISC heterogeneity, with H3K27me3 in differentiated cells and H3K27ac in niche-resident ISCs. Compared to *WT* ISCs, differentiated *WT* villus cells showed dynamic modulation at 3,992 enhancers, associated in aggregate with epithelial differentiation and cell fate commitment genes (Extended Data Fig. 3a). In villus cells, however, only 26% of these sites lost H3K27me3 and gained H3K27ac, indicating acquired enhancer activity; these enhancers lie near genes that serve ion transport, metabolic processes, and other villus functions. Most sites (74%) had diminished H3K27ac in villus cells, while H3K27me3 was preserved or increased, and these sites were linked to genes associated with embryonic development, such as *Sox5*, *Foxa2*, and *Wnt3*. Thus, H3K27me3 at most modulated enhancers marks the stem-cell state. These findings collectively nominate H3K27me3 as a putative gatekeeper of stemness in cells at the crypt base, with progressive loss at *stemness* and *growth* enhancers in *Apc^‒/‒^* cells associated with incursion of stem-like transcriptional programs beyond the usual ISC niche (Fig. 1h).

### Graded activation of silent stemness enhancers during transformation

Negative and positive correlations between H3K27me3 or H3K27ac levels, respectively, and gene expression indicate functional consequences of enhancer modulation. To ask whether the balance of H3K27ac and H3K27me3 across *stemness* and *growth* enhancers reflects quantitative changes in chromatin accessibility, an independent parameter, we profiled mRNA and open chromatin simultaneously in *WT* and *Apc^‒/‒^* transformed epithelial cells at D14 at single-nucleus resolution. Applying stringent quality filters (>12,000 unique molecular identifiers; >1,000 detected genes; mitochondrial ratio <0.35; expression novelty >0.8/cell) to integrated data on gene expression and chromatin accessibility, we identified distinct stem, transit-amplifying (TA), and differentiated cell states (Fig. 2a, Extended Data Fig. 3b). We then derived single-cell activity scores for *stemness* and *growth* enhancers, defined as average enhancer activity scores multiplied by the ATAC-seq signal at each enhancer. In *WT* cells, mean activity scores at *stemness* enhancers were highest in Lgr5^+^ stem cells (0.4) and progressively lower in TA and differentiated (enterocyte) cells; *growth* enhancers, specific to transformed cells, showed little to no accessibility (mean activity scores ≤0.03) in any *WT* cell state (Fig. 2b). In D14 *Apc^‒/‒^*epithelium, by contrast, both stemness and growth enhancers exhibited robust activity across Lgr5⁺ and Lgr5⁻ cells, indicating broad expansion of enhancer activity beyond the stem-cell compartment (Fig. 2b). The concordance between H3K27-defined enhancer states and single-cell chromatin accessibility underscores their activity in ISC differentiation and transformation.

**Figure 2.**
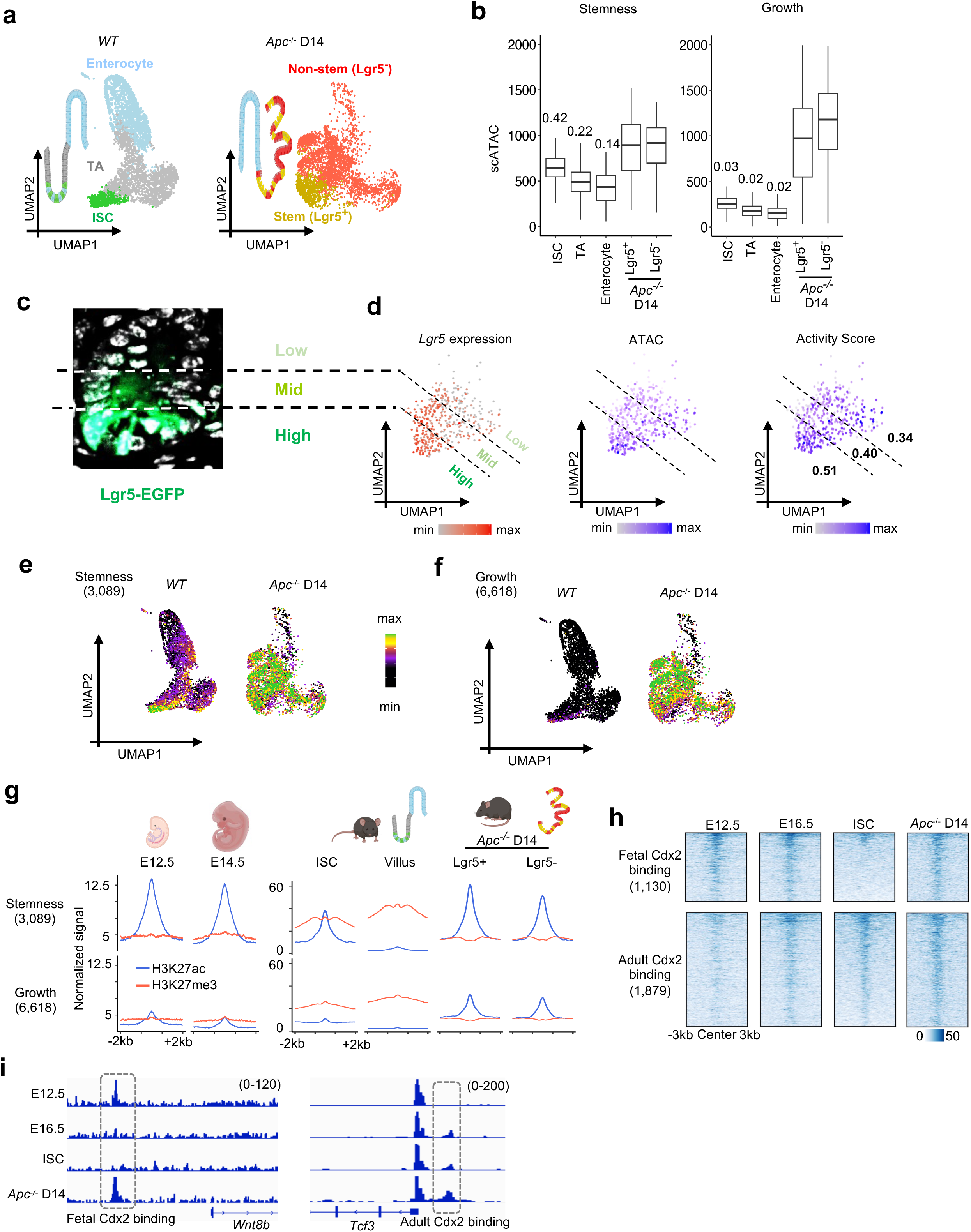
Single-cell multiome analysis reveals activation of stemness and growth enhancers during Apc-deficient transformation at high resolution. **a.** UMAP of scRNA-seq and scATAC-seq from intestinal epithelium showing major epithelial populations in *WT* (ISCs, green; TA, grey; enterocytes, blue) and transformed stem (yellow) and non-stem (red) cells in *Apc^‒/‒^* intestines. **b.** Aggregate chromatin accessibility at stemness and growth enhancers across indicated epithelial populations (mean single-cell enhancer activity score shown above each box). **c.** Representative crypt image showing graded endogenous Lgr5-EGFP expression, highest at the crypt base. **d.** UMAPs of Lgr5 RNA expression, ATAC signal, and enhancer activity score (AS) at 3,089 stemness-linked enhancers within the ISC cluster, demonstrating concordance among these stemness features. **e,f.** UMAP visualization of chromatin accessibility at stemness (**e**) and growth (**f**) enhancers showing ISC-restricted activation of stemness enhancers in WT that expands broadly across *Apc^‒/‒^* cells by day 14, whereas growth enhancers are inactive in *WT* but are robustly activated in transformed cells. **g.** H3K27ac and H3K27me3 signal profiles at stemness (top) and growth (bottom) enhancer sets across fetal intestine (E12.5, E14.5)^26^, adult ISCs, villus cells, and *Apc^‒/‒^* D14 Lgr5^+^ and Lgr5^‒^ populations, showing progressive activation of fetal enhancer programs during transformation. **h.** Heatmaps of Cdx2 occupancy at fetal- and adult-specific binding sites across development (E12.5, E16.5), adult ISCs, and *Apc*^‒/‒^ Lgr5^+^ cells revealing restoration of fetal TF binding in tumours. **i.** Genome browser views illustrating Cdx2 re-engagement at a fetal enhancer near *Wnt8b* in *Apc*^‒/‒^ Lgr5^+^ cells, contrasted with an adult-specific *Tcf3* binding site.

*Lgr5* expression is highest at the crypt base, marks cells with the most stem potential, and decreases in upper crypt tiers, reflecting graded decline of stemness^1^ (Fig. 2c). Within the sc-multiome ISC cluster (Fig. 2a), *WT* cells with high *Lgr5* expression displayed the most chromatin accessibility at *stemness* enhancers and highest activity scores, and both parameters declined in proportion to *Lgr5* mRNA expression (Fig. 2d). Thus, these enhancers are most active in Lgr5^high^ cells and progressively inactivated in Lgr5^low^ and Lgr5^−^ cells by H3K27ac exchange into H3K27me3, thereby restricting stemness to a finite cell fraction. Upon *Apc* loss, both *stemness* and *growth* enhancers were accessible by D14 in both Lgr5^+^ and Lgr5^−^ cells (Fig. 2e,f), revealing spread of an epigenetic mark of stemness into non-stem populations. These findings support the idea that transformed cells progressively activate stemness-associated enhancers by depleting H3K27me3, which normally suppresses stem-like activity in differentiating cells.

### *Stemness* enhancers mark fetal ‘stem’ programs in adult stem cells and fuel their expansion in transformation

Whereas adult intestinal stemness is restricted to Lgr5^+^ cells, the fetal gut epithelium toggles between Lgr5^+^ and Lgr5^−^ states and either population can engender adult ISCs^22^. Analysis of public^23^ and our own data revealed *stemness* enhancer activity in epithelial cells at embryonic days (E) 12.5 and 14.5 (Fig. 2g), with activity scores of 0.72 and 0.60, respectively; in contrast, *growth* enhancers specific to *Apc^‒/‒^* transformed cells were less active at E12.5 (score 0.51) or E14.5 (score 0.34). This H3K27ac dominance over H3K27me3 at *stemness* enhancers in fetal cells resembled that of *Apc^‒/‒^* transformed cells (Fig. 2g). Thus, thousands of enhancers active in early intestine development are retained in a distinct epigenetic state in adult ISCs, are inactive with H3K27me3 marking in differentiated adult cells, and redeploy during transformation. The transcription factor Cdx2 regulates distinct enhancers in fetal and adult intestinal epithelium^24, 25^, and loss of H3K27me3 in adult *Eed^‒/‒^* intestines induces ectopic Cdx2 binding at fetal sites.^25^ CUT&RUN analysis revealed that nearly 40% of *stemness* enhancers coincided with sites that bound Cdx2 at E12.5 and in *Apc^‒/‒^* but not *WT* adult Lgr5^+^ stem cells (Fig. 2h,i). Thus, H3K27me3 represses hundreds of fetal enhancers in adult cells and their reactivation in transformed cells is associated with renewed binding of at least one transcription factor, Cdx2.

Recent studies identify 51 genes, including *Anxa3*, *Krt7*, *Ly*6f and *Col4a2*, that are expressed at higher levels in fetal than in adult intestinal organoids, reactivated during transformation, and implicated in adenoma growth and resistance to cancer therapy^26^. In fetal cells, the promoters of these “onco-fetal” genes carry H3K27ac, but in adult epithelium they also carry H3K27me3, which is lost by D7 after *Apc* deletion (Extended Data Fig. 3c). In contrast, classic stem-cell genes like *Lgr5* and *Smoc2* lack H3K27me3 in ISCs and acquire the mark in differentiated villus cells (Extended Data Fig. 3c-d). Although ∼70% of “oncofetal genes”, e.g., *Anxa3, Anxa5, Samd5* and *Col4a2*, are linked to *stemness* enhancers, *growth* enhancers, or both (Extended Data Fig. 3e), they represent a small fraction of a larger *bona fide* fetal program that shows distinct cis-regulatory states in fetal epithelium and adult ISCs and is reactivated in transformed *Apc^‒/‒^* cells.

### Conserved, H3K27me3-protected stemness is maintained in human CRCs

H3K27me3 prevents fetal gene activity in multiple adult tissues^25, 27^. Although *stemness* and *growth* enhancers were both repressed by H3K27me3 in adult intestine and activated in adenomas, only *stemness* enhancers showed fetal activity (Fig. 2g) and their markedly greater evolutionary conservation compared to *growth* or other intestinal enhancers (Extended Data Fig. 3f) suggests cardinal regulatory functions. Indeed, single-cell RNA-seq data from multiple *APC*-mutant CRC samples^28^ showed that genes linked (±25 kb from TSS) to *stemness* enhancers are expressed in a subset of cells, whereas genes linked to *growth* enhancers are expressed across the population (Fig. 3a). Unsupervised pseudotime analysis^29^ revealed a continuum of cell states that paralleled high-to-low expression of *stemness*–linked genes (Fig. 3a,b) and placed cells with high expression of *stemness*–linked genes on a distinct trunk, while *growth*–linked genes were distributed across trajectories (Fig. 3c,d). Genes linked to *stemness* enhancers, e.g., *ANXA1, MMP7, WNT11,* and *NKD1*, declined along pseudotime, while those linked to *growth* enhancers, e.g., *MYC, CCND1/2,* and *FOS*, were uniformly high (Fig. 3e, Extended Data Fig. 3g). These findings suggest an enhancer-based hierarchy in human CRCs, with H3K27me3-controlled genes active in a restricted truncal (clonogenic) population that seeds non-stem cell trajectories.

**Figure 3.**
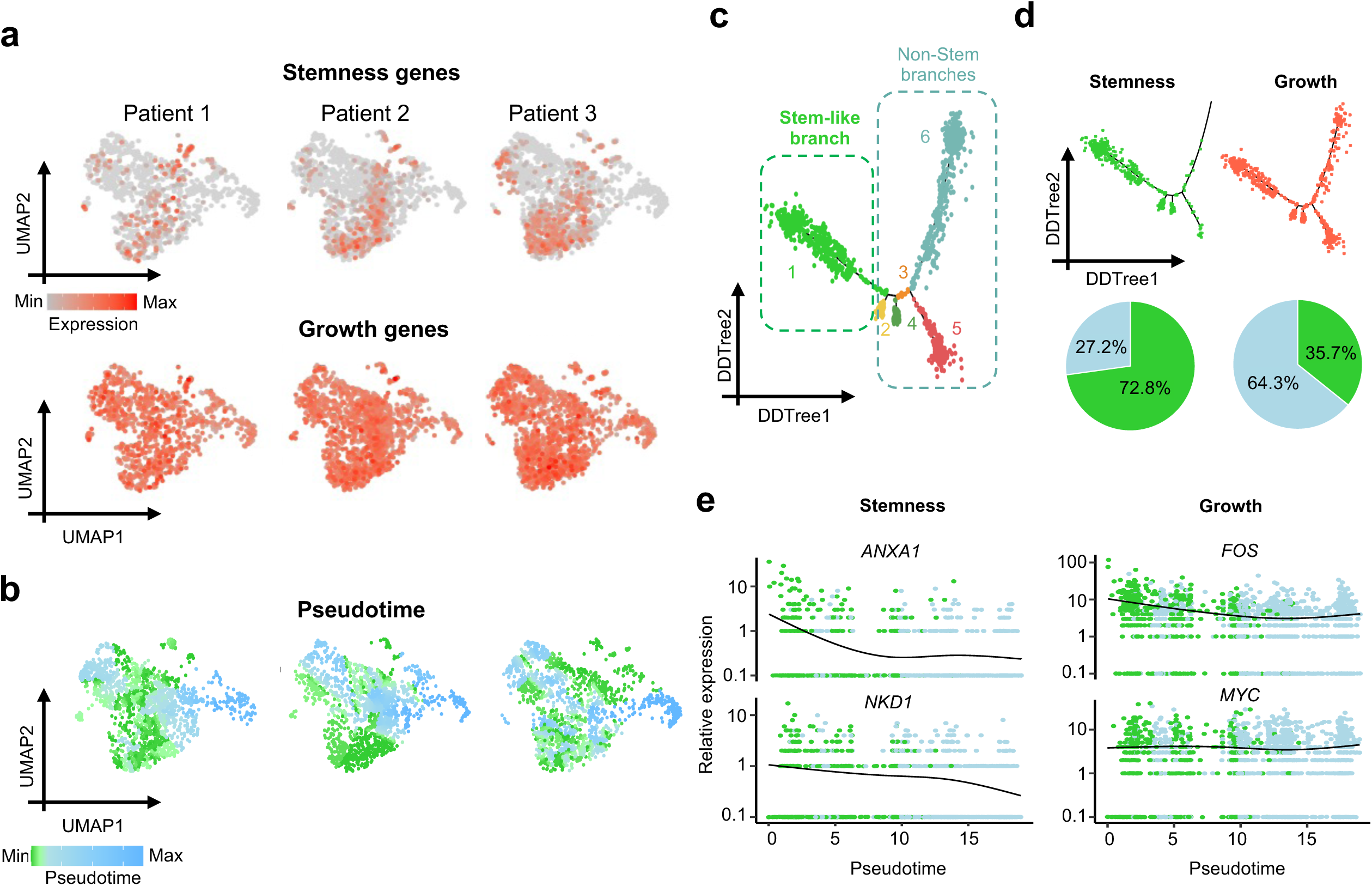
Conserved and H3K27me3-protected fetal regulatory programs are reactivated in transforming cells and sustain activity in advanced colorectal tumours. **a.** UMAP of single-cell transcriptomes from three CRC patients^28^ showing collective expression of stemness- and growth-associated gene signatures. **b.** UMAPs showing pseudotime projections of human tumour cells, inferring differentiation trajectories from stem-like to more differentiated states. **c.** Diffusion map (DDTree) from pseudotime analysis showing differentiation trajectories in human tumours, highlighting stem-like and non-stem branches. **d.** DDTree map of human CRC cells stratified by stemness vs. growth gene signatures identified from *Apc^‒/‒^* mouse adenomas, demonstrating early epigenetic rewiring maintains a minority of stem-like cell population, whereas growth programs remain broadly active in advanced tumours. Pie charts quantify proportions of cells from stem-like (green) and non-stem (blue) branches identified in human tumours. **e.** Expression of stemness (*ANKA1*, *NKD1*) and growth (*FOS*, *MYC*) enhancer linked genes plotted along pseudotime in stem-like (green) and non-stem (blue) cells. Stemness-linked gene expression declines with differentiation, whereas growth gene expression is sustained across most tumour cells. The disparity in cell numbers is expected, as stem-like cells constitute a smaller fraction of the tumour, while most cells express growth genes.

Additionally, TFs such as *Emp1, Prox1,* and *Mex3a* are linked to therapy resistance and relapse,^26, 30–35^ yet they exhibit H3K27me3 erosion and activation across both Lgr5^+^ and Lgr5^−^cells early in cell transformation driven by *Apc* loss (Extended Data Fig. 3h), highlighting the far-reaching impact of early H3K27me3 erosion in CRC.

### H3K27me3 erosion directs superenhancer DNA demethylation in transformed cells

Beyond the 9,707 *stemness* and *growth* enhancers that lost H3K27me3 (Fig. 1f), another 7,798 sites gained H3K27ac in *Apc^‒/‒^* cells at D14 (*q* <0.05, ≥2-fold, Fig. 4a). Notably, 65% of these sites clustered in dense hubs that averaged more than six newly active or pre-existing enhancers within 12.5-kb windows (Extended Data Fig. 4a), resembling superenhancers (SEs)^36, 37^. Separately, the ROSE algorithm^37^ identified 1,342 SEs in D14 *Apc^‒/‒^* stem cells and many newly activated, H3K27me3-independent enhancers contributed to these SEs: 1,214 sites lacked H3K27ac in *WT* cells (*de novo*), while 2,779 loci with previous H3K27ac signal were strongly increased (Fig. 4b-d). Some *de novo* enhancers arose within the boundaries of 994 pre-existing SEs in *WT* ISCs, while others expanded beyond them, increasing the median of 4 constituent enhancers per SE to 10 (Fig. 4c-e). *Stemness* and *growth* enhancers were hypomethylated (≤50% methylated CpG residues) (Extended Data Fig. 4b). In contrast, although *de novo* enhancers lacked H3K27me3 in *WT* or *Apc^‒/‒^* cells, 92% showed significant DNA hypomethylation (*p* <0.001, Δ≥40%, Fig. 4b,d), revealing H3K27me3-independent DNA demethylation and SE activation.

**Figure 4.**
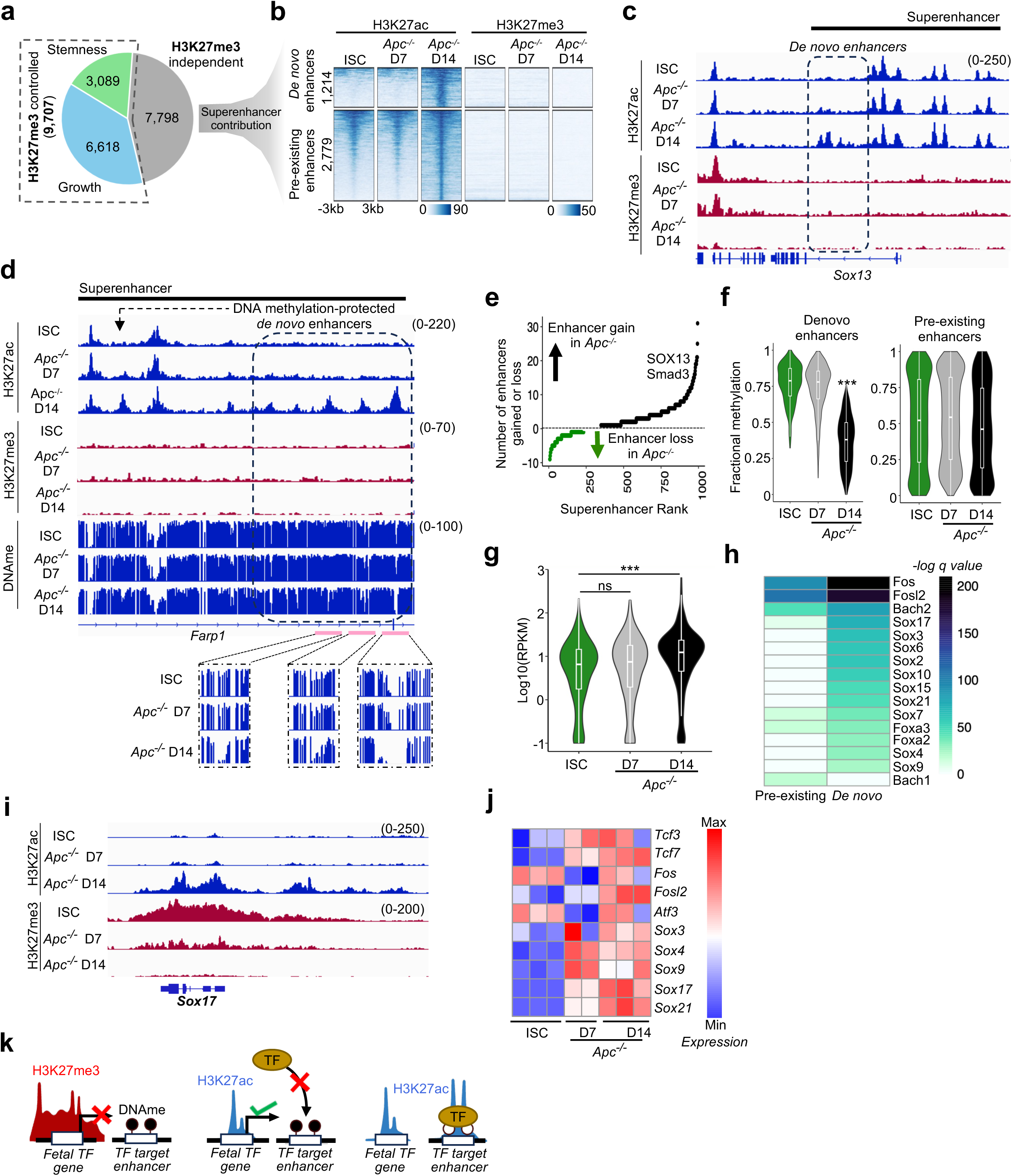
H3K27me3 erosion precedes and directs DNA methylation changes, leading to superenhancer remodeling during cell transformation. **a.** Pie-chart showing H3K27me3 controlled (stemness and growth) enhancers and H3K27me3 independent enhancers activated in *Apc^‒/‒^* Lgr5⁺ cells at D14. Many H3K27me3 independent enhancers contribute to superenhancers. **b.** Heatmaps of H3K27ac and H3K27me3 signals at *de novo* and pre-existing enhancers formed within superenhancers in *Apc^‒/‒^* day 14 cells. **c.** Genome browser tracks of H3K27ac and H3K27me3 signals showing *de novo* enhancers contributing to a superenhancer near *Sox13*. **d.** Genome browser views showing H3K27ac enrichment and hypomethylation at *de novo* methylation-protected enhancers within a superenhancer near *Farp1* locus in *Apc^‒/‒^* Lgr5^+^ cells. **e.** Scatter plot showing gains or loses of enhancers within superenhancers during ISC transformation into *Apc^‒/‒^* Lgr5⁺ cells at D14, demonstrating an increase in the number of enhancers in transformed stem cells. **f.** Violin plots showing the loss of DNA methylation at *de novo* enhancers, but not at pre-existing enhancers, in *Apc^‒/‒^*D14 cells. **g.** Expression of genes associated with newly formed superenhancers in *Apc^‒/‒^* D14 cells (***, p < 0.001, one-way Wilcoxon). **h.** Heatmap showing transcription factor motif enrichment at *de novo* and pre-existing enhancers within superenhancer domains. **i.** Genome browser plots showing rapid loss of H3K27me3 at Sox17 locus allowing activation of multiple enhancers at that locus during cell transformation. **j.** Heatmap showing gene expression changes for transcription factors with motif enrichment within *de novo* and pre-existing enhancers across ISCs and *Apc^‒/‒^*D7 and D14 stem cells. **k.** Model illustrating that erosion of H3K27me3 marks precedes DNA demethylation, activates fetal TFs that bind to target enhances leading to demethylation and activation of enhancers that drive stem cell transformation and tumour morphological progression.

Whole genome bisulfite sequencing (WGBS) of *Apc^‒/‒^* Lgr5^+^ stem cells further revealed that SE enhancers with *de novo* DNA demethylation at D14 were methylated at D7 (Fig. 4d, f). Moreover, compared to *WT* ISCs, genes associated with these SEs were upregulated only at D14, not at D7 (Fig. 4g); Gene Ontology implicated these genes in cell proliferation, motility, and cell-cell adhesion (Extended Data Fig. 4c), e.g. *Farp1*, a known gastric cancer proliferation factor^38^ (Fig. 4d). Thus, DNA demethylation follows initial H3K27me3 depletion and activation of *stemness* enhancers, marking the second phase of enhancer reprogramming associated with *Apc^‒/‒^*tumourigenesis. These enhancers were highly methylated during embryonic development, as far back as E6.5 endoderm (Extended Data Fig. 4d), indicating that their activation is tumour-specific rather than a recapitulated fetal program. While SEs are typically CpG-rich and hypomethylated^39^, our findings identify subsets of SE sites that are inactive owing to DNA methylation in normal intestinal cells and selectively activated in transformation.

Focal DNA demethylation at enhancers arises from TF binding.^25, 40^ Motif analysis by HOMER^41^ predicted that pre-existing SE enhancers preferentially bind Jun/Fos/AP-1 factors, whereas *de novo* enhancers following *Apc* loss are enriched for Sox-family TF motifs (Fig. 4h). Indeed, Sox17 and other factors active in fetal endoderm but silent in adult intestine are aberrantly reactivated in early murine and human CRC^26, 42^. Among genes associated with *stemness* and *growth* enhancers, 1,132 also showed ≥2-fold promoter H3K27me3 loss (q <0.05) in *Apc^‒/‒^* cells; these genes were highly enriched for fetal programs (GO analysis, *q* <0.001) and encompassed 108 TFs (Extended Data Fig. 4e,f), including *Sox17*, whose broad H3K27me3 repression in *WT* was lost progressively in *Apc^‒/‒^* cells (Fig. 4i). mRNA-seq confirmed expression of *Sox17* and other Sox factors in *Apc^‒/‒^*cells by D7 (Fig. 4j), indicating that early H3K27me3 depletion reactivates fetal TFs, which in turn promote focal DNA demethylation and enhancer activation during transformation.

Notably, native-ChIP-seq for H3K27ac in *Apc^‒/‒^* non-stem (Lgr5^−^) progeny revealed absence of SE enhancer activation at D7 and presence at D14, whereas WGBS of *Apc^‒/‒^* Lgr5^−^ cells lacked significant DNA methylation changes at D14 (Extended Data Fig. 4d,g). Thus, DNA demethylation follows H3K27me3 erosion and enhancer activity gained in Lgr5⁺ stem cells persists in their progeny. Together, these findings define a stepwise cascade, with early H3K27me3 erosion activating plasticity and growth enhancers and fetal TFs, which subsequently drive DNA demethylation and SE remodeling to promote tumour progression. Thus, sequential losses in H3K27me3 and methylated DNA jointly underlie enhancer activity and tumourigenesis (Fig. 4k).

### Promoting H3K27me3 loss accelerates adenoma progression

Early H3K27me3 loss and associated expansion of stemness during transformation suggest a tumour-suppressive role for Polycomb Repressive Complex 2 (PRC2), which deposits H3K27me3. To test how PRC2 modulation influences the stemness–differentiation imbalance in adenomas, we generated *Apc^Fl/Fl^*;*Eed^Fl/Fl^*;*Lgr5^EGFP-CreER-T2^;Rosa26^LsL-tdTomato^*(hereafter *Apc^‒/‒^ Eed^‒/‒^*) mice, in which PRC2 activity is abrogated through Cre-mediated deletion of *Eed*, an essential factor for PRC2 stability and H3K27me3 deposition,^27, 43^ alongside *Apc* loss (Fig. 5a). Efficient *Eed* deletion was confirmed in both Lgr5^+^ and Lgr5^‒^ compartments by PCR on FACS-sorted cells (Extended Data Fig. 5a) and H3K27me3, present in normal epithelium, was absent in adjacent *Apc^‒/‒^ Eed^‒/‒^*tumours (Extended Data Fig. 5b). Conversely, to augment H3K27me3 deposition during transformation, we generated *Apc^Fl/Fl^*;*Col1a*^LSL-Ezh2^;*Lgr5^EGFP-CreER-T2^;Rosa26^LsL-tdTomato^* (hereafter *Apc^‒/‒^ Ezh2^OE^*) mice, in which Cre-mediated overexpression of human *EZH2*, the PRC2 catalytic subunit, elevates H3K27me3 levels (Fig. 5a).^44^ Human *EZH2* transcripts were detected in both Lgr5^+^ and Lgr5^‒^ epithelial populations of *Apc^‒/‒^ Ezh2^OE^* tumours (Extended Data Fig. 5c). We hypothesized that reciprocal perturbations of the H3K27me3 epigenetic barrier would substantially affect adenoma progression and tissue morphology (Fig. 5a).

**Figure 5.**
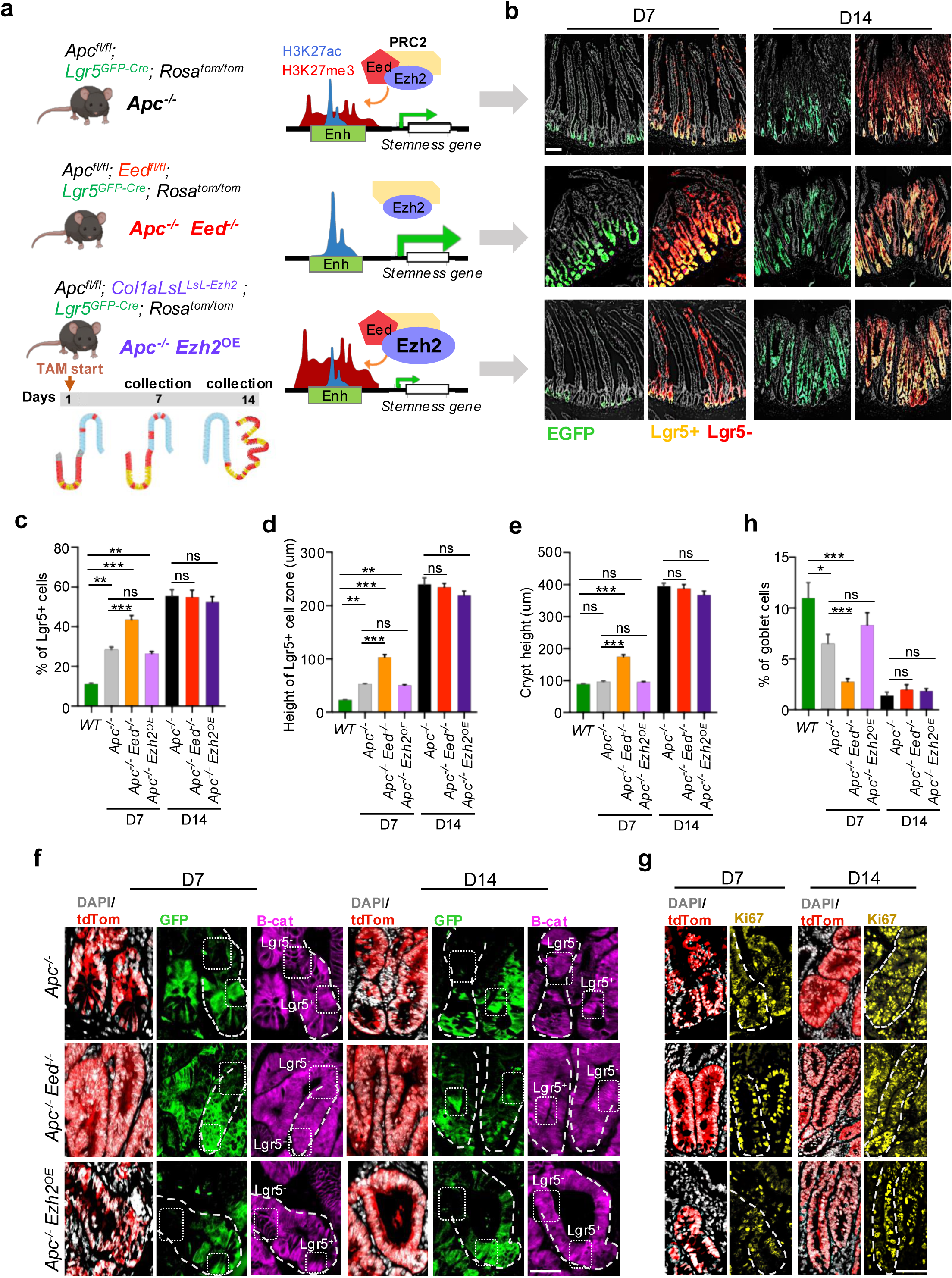
Genetic alteration of PRC2 action influences cell transformation and adenoma growth. **a.** Mouse models for facilitated H3K27me3 loss (*Eed^‒/‒^*) or deposition (*Ezh2^OE^*) alongside *Apc^‒/‒^.* Schematic showing that loss of PRC2 in *Apc^‒/‒^ Eed^−/–^* mice amplifies stemness gene expression, whereas Ezh2 overexpression in *Apc^‒/‒^ Ezh2^OE^* mice suppresses these genes. **b.** Representative immunofluorescence images showing expansion of transformed Lgr5^+^ stem cells (yellow) and emergence of Lgr5^‒^ non-stem cells (red) in *Apc*^‒/‒^, *Apc*^‒/‒^ *Eed*^‒/‒^, and *Apc*^‒/‒^*Ezh2*^OE^ intestines at D7 and D14. *Apc*^‒/‒^ *Eed*^‒/‒^ adenomas display extensive Lgr5^+^ expansion by D7, resembling the D14 architecture of *Apc*^‒/‒^intestine. Scale bar = 100um. **c–e.** Quantification of morphological changes in various adenoma models **(c)** percentage of Lgr5⁺ cells among all transformed (tdTom^+^) cells, **(d)** extent of Lgr5^+^ cell expansion along the crypt–villus axis, and **(e)** crypt height among all transformed cells, highlighting significantly increased Lgr5^+^ cell abundance and a marked reduction in goblet cells compared with PRC2-intact mice (mean ± s.e.m.; \**p* < 0.05, \*\**p* < 0.01, \*\*\**p* < 0.001, ns = not significant). **f.** Immunofluorescence staining for tdTomato (transformed cells), EGFP (Lgr5^+^ stem cells) and β-catenin at D7 and D14 across tumour types. *Apc^‒/‒^ Eed^−/–^* tumours display increased nuclear β-catenin accumulation in Lgr5^‒^non-stem cells at D7, which is observed in *Apc^‒/‒^* only at D14. Scale bar = 50um. **g.** Immunostaining for MKi67 and tdTomato showing increased proliferation in both Lgr5^+^ and Lgr5^‒^ cells in *Apc^‒/‒^ Eed^−/–^* tumours compared with other models. **h.** Quantification of percentage of goblet (Mucin^+^) cells among all transformed cells.

Tumour progression was indeed accelerated in *Apc^‒/‒^ Eed^‒/‒^* mice, which survived a median of ∼2 days (N >10 mice) less than *Apc^‒/‒^* or *Apc^‒/‒^ Ezh2^OE^* cohorts (Extended Data Fig. 6a). By D7, *Apc^‒/‒^ Eed^‒/‒^* intestines showed marked expansion of Lgr5^+^ stem cells, disruption of crypt–villus architecture, and pronounced crypt enlargement, with adenoma dimensions comparable to D14 *Apc^‒/‒^*adenomas (Fig. 5b-e). At this time, *Apc^‒/‒^Eed^‒/‒^*mice also showed increased accumulation of nuclear β-catenin and elevated proliferation (MKi67 staining) in both Lgr5^+^ and Lgr5^‒^ compartments, resembling D14 *Apc^‒/‒^* adenomas (Fig. 5f-g, *n* = 3). *Apc^‒/‒^Ezh2^OE^* did not show significant change in tumour morphology as compared with D14 *Apc^‒/‒^*adenomas within the defined 14-day experimental timeframe (Fig. 5b-g). Consistent with accelerated expansion of stemness, *Apc^‒/‒^ Eed^‒/‒^* mice at D7 exhibited markedly reduced differentiation, evidenced by fewer *Mucin*^+^ goblet cells and a front of *Alpi⁺* enterocytes shifted farther from the crypt base, indicating an expanded stem-cell zone compared to PRC2-proficient tumours (Fig. 5h; Extended Data Fig. 6b,c). Thus, weakened H3K27me3-mediated gatekeeping of *stemness* and *growth* enhancers accelerates stem-cell expansion, adenoma growth, and host mortality.

### Reducing H3K27me3 accelerates enhancer reprogramming and spread of stemness

To assess whether the speed of morphological changes in *Apc^‒/‒^ Eed^‒/‒^* tumours reflected accelerated activation of *stemness* and *growth* enhancers, we profiled H3K27ac and H3K27me3 in both mouse models 7 and 14 days after Cre induction. Compared to *Apc^‒/‒^* stem cells, *Apc^‒/‒^Eed^‒/‒^* stem cells showed faster H3K27me3 erosion at both *stemness* and *growth* enhancers (Fig. 6a), accompanied by increased enhancer H3K27ac (Fig. 6b, Extended Data Fig. 7a). By D7, reduced H3K27me3 and elevated H3K27ac levels were already comparable to those in D14 *Apc^‒/‒^* stem cells, indicating faster engagement of both enhancer classes, as seen in cumulative profile plots (Fig. 6b-c) and quantified using the Kolmogorov-Smirnov (KS) test (Extended Data Fig. 7b–e). Conversely, enhancing PRC2 activity by EZH2 overexpression slowed H3K27me3 loss in *Apc^‒/‒^ Ezh2OE* adenomas, delaying their activation (Fig. 6b-d). In line with these findings, genes near *stemness* and *growth* enhancers were significantly upregulated in *Apc^‒/‒^ Eed^‒/‒^* adenomas at D14 compared to *Apc^‒/‒^* or *Apc^‒/‒^ Ezh2OE* (Fig. 6d). Accelerated H3K27me3 loss at these enhancers in *Apc^‒/‒^ Eed^‒/‒^* adenomas drove mRNA levels at D7 to those seen in *Apc^‒/‒^*adenomas at D14. Thus, PRC2-mediated repression is a tumour-suppressive barrier; its loss enables transcriptional programs that expand stemness, promote adenoma growth, and reduce host survival.

**Figure 6.**
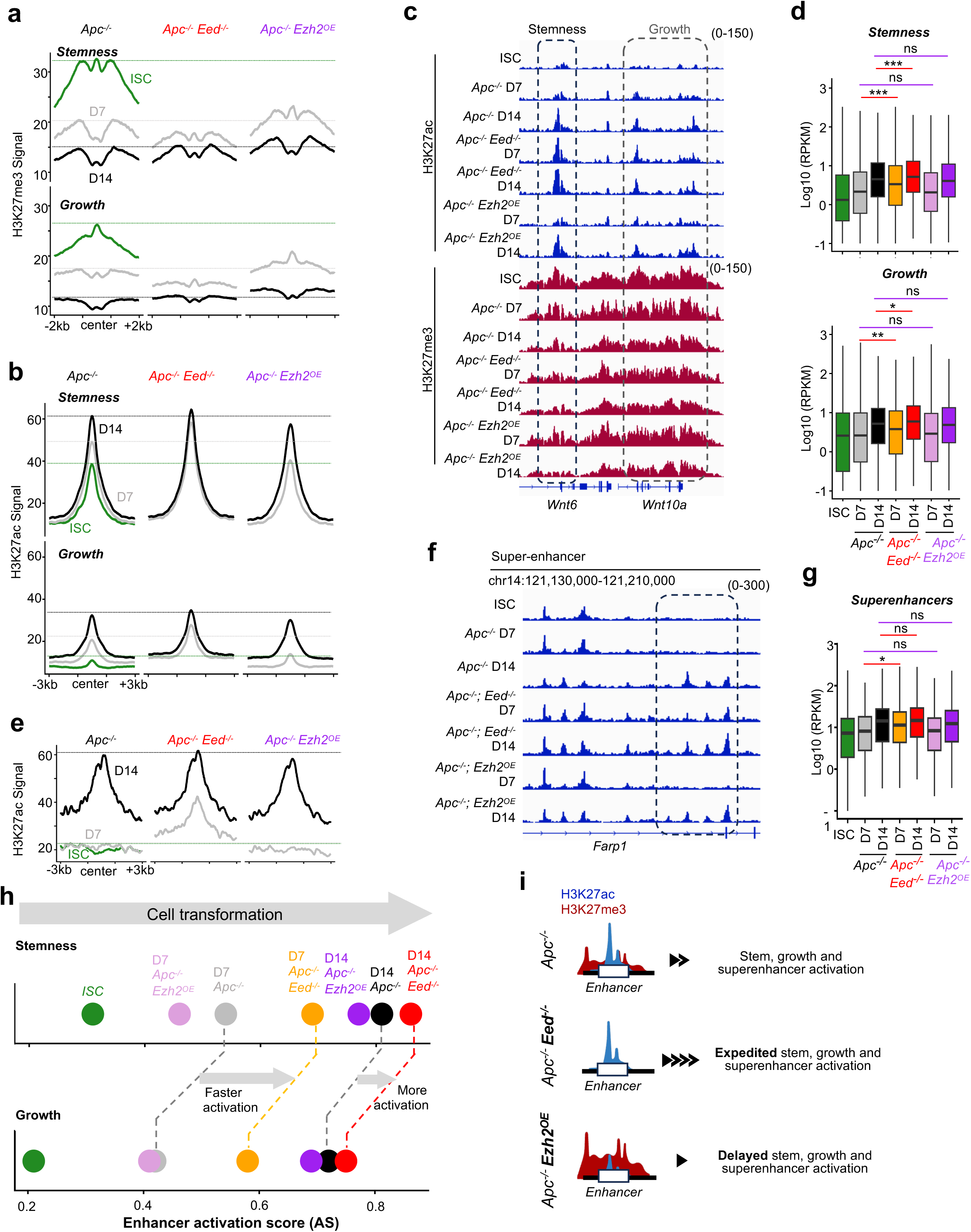
PRC2 modulation alters enhancer reprogramming and cell transformation. **a,b.** Profile plots showing normalized H3K27me3 (a) and H3K27ac (b) signals at stemness and growth enhancers across ISCs and Lgr5^+^ populations from D7 and D14 *Apc^‒/‒^*, *Apc^‒/‒^ Eed^‒/‒^* and *Apc^‒/‒^ Ezh2*^OE^ epithelium. Green, grey, and black dotted lines indicate the highest signal intensity in ISC, D7 *Apc^‒/‒^*, and D14 *Apc^‒/‒^*, respectively. **c.** Genome browser tracks showing H3K27ac and H3K27me3 signal in Lgr5^+^ cells from various mouse models showing accelerated activation of growth enhancers in *Apc^‒/‒^ Eed^‒/‒^* at D7. **d.** Expression of genes uniquely associated with stemness and growth enhancers show significantly faster (D7) and higher activation of linked genes (D7 and D14) in *Apc^‒/‒^ Eed^‒/‒^* as compared to the other two genotypes (*Apc^‒/‒^* and *Apc^‒/‒^ Ezh2*^OE^) (Box and whiskers represent Q1,median, and Q3; \**p* < 0.05, \*\**p* < 0.01, \*\*\**p* < 0.001, ns = not significant). **e.** Profile plot of H3K27ac signals at *de novo* enhancers within superenhancers in Lgr5^+^ cell populations from D7 and D14 *Apc^‒/‒^*, *Apc^‒/‒^ Eed^‒/‒^* and *Apc^‒/‒^Ezh2*^OE^ mice. Green, grey, and black dotted lines mark maximal signal intensities in ISC, D7 *Apc^‒/‒^*, and D14 *Apc^‒/‒^* cells, respectively. **f.** Genome browser tracks showing H3K27ac signal across Lgr5^+^ cell population across indicated samples near *Farp1* locus. Dotted box highlights multiple *de novo* enhancers activated early at D7 only in *Apc^‒/‒^ Eed^‒/‒^* cells. **g.** Genes associated with *de novo* enhancers within superenhancers show significant early (D7) gain in expression only in *Apc^‒/‒^ Eed^‒/‒^* cells. **h.** Dot plots showing average enhancer activation scores for stemness and growth enhancers for various genotypes at D7 and D14 of transformation. *Apc^‒/‒^ Eed^‒/‒^* cells exhibit faster as well as more activation of both types of enhancers as compared to *Apc^‒/‒^* cells. **i.** Model illustrating expedited epigenomic remodeling upon PRC2 inactivation through lowering of H3K27me3 and DNA hypomethylation, facilitating ISC transformation and adenoma growth.

SE expansion, typically restricted to D14 *Apc^‒/‒^* transformed cells, was also evident in *Apc^‒/‒^ Eed^‒/‒^* tissues at D7 (Fig. 6e-g; Extended Data Fig. 7f,g), accompanied by precocious induction of SE-associated genes (Fig. 6g). Enhancer activation scoring (Fig. 1e) revealed that beyond accelerated remodeling, PRC2 loss (*Apc^‒/‒^ Eed^‒/‒^*) drove H3K27ac levels at *stemness* and *growth* enhancers higher than those observed in *Apc^‒/‒^* cells at D7 (Fig. 6h). Uniscale reduction analysis using the MASS algorithm supported these observations (Extended Data Fig. 7h). Thus, H3K27me3 loss accelerates enhancer activation, transcriptional reprogramming, and emergence of adenomas, establishing PRC2 as a safeguard of stem-cell enhancer activity (Fig. 6i).

### Disruption of H3K27me3 amplifies stemness and fitness of transformed cells

Transformed stem cells fuel tumour growth and the capacity of non-stem tumour cells to regain stemness allows tumours to regenerate after the stem-cell compartment is eradicated^15^. Emergence and expansion of aggressive clones through clonal competition can further advance tumour persistence and growth. While mutational and microenvironmental drivers of stemness and fitness have been well studied,^45^ the expanded stem-like pool in *Apc^‒/‒^ Eed^‒/‒^* adenomas suggests that accelerated activation of *stemness* and *growth* enhancers might confer a competitive growth advantage.

To test expanded stemness functionally, we evaluated organoids derived from FACS-purified Lgr5^+^ and Lgr5^‒^ cells at D7 and D14 across PRC2 genotypes. As expected, both populations from D14 tumours formed organoids at all densities from 1,000 to 10,000 cells per dome (Fig. 7a,b). Strikingly, Lgr5^‒^ cells from *Apc^‒/‒^ Eed^‒/‒^* mice at D7 formed organoids when seeded at high density, indicating that rapid H3K27me3 loss enables the spread of stem-like potential beyond the Lgr5^+^ compartment. To determine whether this spread reflects accelerated propagation of enhancer rewiring in Lgr5^‒^ progeny, we used native-ChIP-seq to profile H3K27ac gain and H3K27me3 erosion at *stemness* and *growth* enhancers and SEs in Lgr5^‒^ cells across all genotypes at D7 and D14 (Fig. 7c-e). Only Lgr5^‒^ cells from *Apc^‒/‒^ Eed^‒/‒^*mice gave strong H3K27ac signals all three enhancer classes by D7, linking rapid enhancer remodeling to accelerated spread of stemness.

**Figure 7.**
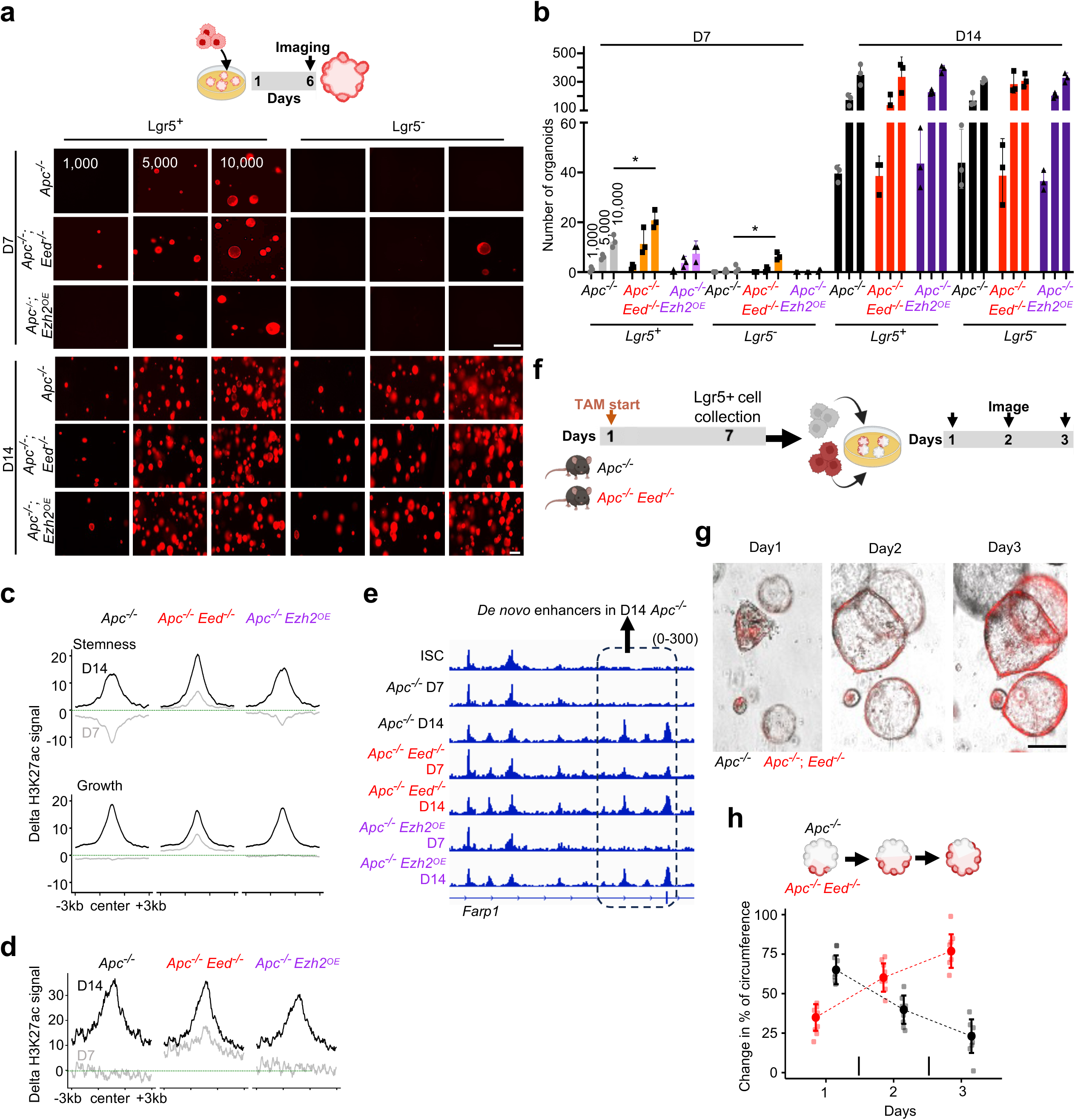
H3K27me3 barrier disruption promotes stemness and competitive fitness during transformation. **a.** Schematic illustrating the experimental design for the organoid formation assay using FACS-sorted Lgr5^+^ and Lgr5^‒^ populations from D7 and D14 tumours across indicated genotypes. Representative fluorescence images show organoids formed at day 6 from 1,000, 5,000, and 10,000 initial cells derived from Lgr5^+^ and Lgr5^‒^ populations. Scale bar = 50um. **b.** Quantification of organoid formation at day 6 shows that only *Apc^‒/‒^ Eed^‒/‒^* cells at D7 have transformed stem cell potential allowing organoid growth in the absence of Wnt (*p* < 0.05). **c.** Profile plots showing differential H3K27ac signal between Lgr5^‒^ tumour samples and ISCs at stemness and growth enhancers. **d.** Profile plots showing differential H3K27ac signal between Lgr5^‒^ tumour samples and ISCs at *de novo* enhancers formed within *Apc^‒/‒^* superenhancers. **e.** Genome browser tracks showing H3K27ac signal in Lgr5^‒^ non-stem cells across indicated samples near the *Farp1* locus, highlighting strong activation of *de novo* enhancers in *Apc^‒/‒^ Eed^‒/‒^* cells at D7. **f.** Schematic of the competitive hybrid organoid assay. **g.** Overlay of brightfield and fluorescence images showing hybrid organoids composed of *Apc^‒/‒^ Eed^‒/‒^*(tdTomato^+^, red) and *Apc^‒/‒^* (unlabeled) cells, demonstrating competitive dominance of *Apc^‒/‒^ Eed^‒/‒^* cells. **h.** Quantification of the relative contribution of *Apc^‒/‒^* (unlabeled) and *Apc^‒/‒^ Eed^‒/‒^* (red) cells to the total organoid circumference over three days of co-culture.

To test whether this enhancer reprogramming confers a competitive advantage, we performed clonal-competition assays. We co-cultured 50,000 unlabeled Lgr5^+^ adenoma cells from *Apc^‒/‒^* mice at D7 with an equal number of tdTomato-labeled cells from *Apc^‒/‒^ Eed^‒/‒^* mice (Fig. 7f) and tracked relative clone dimensions over 3 days (Fig. 7g and Extended Data Fig. 7i). *Apc^‒/‒^Eed^‒/‒^* clones rapidly outcompeted *Apc^‒/‒^* clones, dominating hybrid organoids by day 3 (Fig. 7g,h) and revealing a fitness advantage from accelerated enhancer activation. Thus, enhancer remodeling initiated by H3K27me3 erosion underlies *Apc^‒/‒-^*transformation and accelerating this erosion amplifies stem cell potential, suppresses differentiation, and promotes clonal dominance. H3K27me3 gatekeeping regulates the pace of tumour initiation and also enhances fitness.

## DISCUSSION

Most cells in the fetal intestinal epithelium can give rise to adult stem cells, suggesting that developmental cell plasticity becomes restricted to a select cell population in adults.^2, 3^ Although extrinsic signals from the crypt niche, particularly Wnt/Rspondin, are known to regulate stemness, cell-intrinsic controls that limit stemness to the crypt base are poorly understood. The tissue microenvironment also influences tumour development and growth, but expanded stemness and cell plasticity precede the known external cues^1, 33, 46^, underscoring the importance of understanding cell-intrinsic mechanisms that contain homeostatic plasticity and enable its expansion in tumours. Epigenetic barriers have long been proposed as intrinsic gatekeepers of tumourigenesis,^13^ yet their molecular identity remains elusive. Our findings provide a unifying molecular model for homeostasis and transformation: Stemness-related gene activity in fetal epithelium supports cell plasticity and rapid tissue growth, whereas in adult intestine, these genes and their enhancers are selectively active in stem cells; enhancer H3K27me3 represses these programs in non-stem cells, hence compartmentalizing stemness and cell differentiation. Loss of this epigenetic safeguard dismantles constraints, enabling cell transformation. Further emphasizing the critical role of this intrinsic control, even *Apc*-mutant cells, which acquire Wnt-independent self-renewal, continue to generate differentiated progeny. Thus, a cell-intrinsic epigenetic modification that demarcates stemness from differentiated cells is eroded in concert with external cues during transformation. Our detailed temporal dissection of this process resolved sequential steps, distinguishing early expansion of stemness from later compromise in differentiation. Together, these findings reveal stepwise, cell-intrinsic epigenetic remodeling that dismantles homeostatic control of stemness to drive tumourigenesis.

Fetal gene expression in diverse CRC models and patient tumours^26, 30–33^ points to ectopic reactivation of developmental transcription. Our findings reveal that fetal gene expression in intestinal tumours does not represent fortuitous activity of selected fetal genes, but a coordinated re-deployment of stemness-related developmental programs. They show how H3K27me3 restricts these fetal signatures in normal adult intestinal epithelium and how they are reactivated in two successive phases during tumourigenesis. Sequential activation of these genes aligns with evolving tumour tissue remodeling, mirroring how these programs are deployed during fetal life. Importantly, even beyond transformation, fetal genes reactivated by H3K27me3 erosion remain expressed in advanced and metastatic tumours.

Metastatic clones are known to arise early, before detectable tumour formation,^47^ and their mutational profiles are largely the same as their primary sources,^48^ implying that metastatic capabilities arise from early molecular reprogramming. Early erosion of H3K27me3 at the onset of cell transformation may explain this early reprogramming, illustrated by H3K27me3 loss near *Emp1, Prox1, Mex3a,*^8–11^, and other genes associated with drug resistance and disease relapse within 14 days of *Apc* inactivation. Interestingly, while most metastatic cells lack *Lgr5* expression,^8–12^ they regenerate Lgr5^+^ populations at secondary sites, enabling survival and tumour growth.^8–11^ Moreover, ablation of Lgr5^+^ cells by radiation or chemicals leads to tumour regression, yet Lgr5⁻ cells swiftly restore Lgr5^+^ populations once treatment ceases^26, 30–35^. Therefore, both metastasis and post-therapy relapse reflect plasticity between cell types. Our data show that loss of enhancer H3K27me3 equalizes Lgr5^+^ and Lgr5^‒^ chromatin states, associated with dynamic interconversion between these distinct cell types.

Suppressing cells with active fetal and relapse potential is proposed as a therapeutic strategy to prevent disease recurrence^8–12^; this makes it important to understand the epigenetic basis of distinct transcriptional programs. Our investigation of Lgr5^+^ and Lgr5⁻ populations at different steps in transformation identified enhancer H3K27me3 as a key gatekeeper in adult tissues and its erosion as a central event that first enables expansion of Lgr5^+^ tumour stem cells and, subsequently, molecular equivalence between transformed Lgr5^+^ stem cells and their Lgr5⁻ progeny that facilitate disease relapse. Together, these insights establish a unifying framework for the regulation of stemness from fetal development to adult tissue homeostasis and tumourigenesis, laying the foundation for strategies to predict and contain stemness and cell plasticity.

## METHODS

### EXPERIMENTAL MODEL AND SUBJECT DETAILS

#### Mouse Models

All mice were handled and maintained following the IACUC protocol approved by the Department of Animal Resources at the University of Southern California. Animals were housed under 12-hour light/dark cycles, at 23 ± 1°C, with 50 ± 10% humidity. Mice displaying hunched posture, ruffled fur, decreased movement, or a rapid 20% body weight loss were removed from the study and humanely euthanized. Euthanasia was conducted via CO2 inhalation for five minutes, using a flow rate approximating 20% of chamber volume per minute. Cervical dislocation was applied as a secondary euthanasia method. *ROSA26^LSL-tdTomato^* (strain 007909) and *Lgr5^EGFP-CreER^* (strain 008875) mice were purchased from The Jackson Laboratories; *Apc^fl^* mice were a gift from C. Perret,^49^ and *Col1a^LSL-Ezh2^* mice were a gift from K Wong.^44^ Genotyping by PCR confirmed the presence of genetically modified alleles both at weaning and during experiments (Extended Data Fig. 5a). To induce deletion of *Apc* and *Eed*, and expression of *EZH2*, adult male and female mice aged eight weeks or older received 2 mg tamoxifen via intraperitoneal injection on two consecutive days. To prevent escape from Cre-mediated excision^19^, additional tamoxifen injections of 1 mg were administered every third day. Tissues were harvested at the designated time points for each experiment.

#### Isolation of intestinal epithelial cells

Adult male and female mice of age 8 weeks or older were used for isolation of intestinal epithelial cells from the proximal one-third of the small intestine. After dissection, the intestine was opened lengthwise exposing the epithelium and rinsed with cold phosphate-buffered saline (PBS) multiple times. The tissue was then placed in 5 mM EDTA in PBS (pH 8.0) at 4 °C for 30 minutes, with vigorous shaking every 10 minutes to aid dissociation. Crypts and villi were isolated by centrifugation at 400 × g for 5 minutes and incubation in 20 ml of 4% TrypLE (Invitrogen) in DMEM at 37 °C for 20 minutes with rotation to generate single-cell suspensions. After stopping the reaction using of 15 ml DMEM, cells were pelleted by centrifugation. For ISC isolation, fluorescence-activated cell sorting (FACS) was conducted using a BD SORP FACSymphony S6 instrument to capture viable (DAPI−) GFP+ cells based on endogenous GFP expression (Extended Data Fig. 1j). Apc-null *Lgr5^+^* and *Lgr5^−^* cells were isolated in a similar fashion using endogenous GFP and tdTomato signals.

#### Intestinal organoid generation

Crypts were isolated from the proximal one-third of the small intestine of adult male and female mice aged 8 weeks or older. Immediately following dissection, the intestine was opened longitudinally, rinsed with cold phosphate-buffered saline (PBS), and incubated in 5 mM EDTA in PBS (pH 8.0) at 4 °C for 15 minutes with rotation. Tissue fragments were pelleted at 300 × g for 5 minutes, the supernatant was removed, and fresh 5 mM EDTA was added. The tissue was incubated on a rocker at 4 °C for an additional 25 minutes to release crypts. The dissociated epithelium was passed through a 70 μm cell strainer, and the flow-through containing crypts was collected and centrifuged at 400 × g for 5 minutes at 4 °C. The crypt pellet was washed three times with ice-cold Advanced DMEM/F12 (Gibco) and pelleted each time at 400 × g for 5 minutes at 4 °C. Approximately 2 μl of the final crypt pellet was resuspended in 50 μl of Matrigel (Corning, 356231) and plated as 50 μl domes in 24-well flat-bottom tissue culture plates. Domes were polymerized at 37 °C for 10 minutes before adding culture medium. Organoids were maintained in mouse intestinal organoid media composed of Advanced DMEM/F12 (Gibco, 12634-028) supplemented with 10% R-spondin-conditioned medium, 1× GlutaMAX (Gibco, 35050-061), 10 mM HEPES (Gibco, 15630-080), 0.2% Primocin (Invitrogen, ant-pm-2), 0.2% Normocin (Invitrogen, ant-nr-2), 1% B-27 (Gibco, 12587-010), 0.5% N-2 (Gibco, 17502-048), 1.25 mM N-acetylcysteine (Sigma, A8199), 50 ng ml⁻¹ EGF (Peprotech, 315-09), 100 ng ml⁻¹ Noggin (Peprotech, 250-38), and 10 μM Y-27632 (Sigma, Y0503). Organoids were cultured at 37 °C in a humidified incubator with 5% CO₂, and media was changed every 3 days. Organoids were passaged as needed or every week.

#### Organoid formation assay for stemness

A total of 1,000, 5,000, or 10,000 FACS-sorted Lgr5^+^ and Lgr5^‒^ epithelial cells were isolated from day 7 and day 14 *Apc^‒/‒^*, *Apc^‒/‒^ Eed^‒/‒^*, and *Apc^‒/‒^Ezh2^OE^* intestinal tissues and embedded in 30 μl of Matrigel (Corning, 356231) per well. Embedded cells were plated in 24-well plates and cultured for 6 days in mouse intestinal organoid medium (see above) lacking R-spondin-conditioned medium and Y-27632. Organoids were imaged on day 6 using a Keyence BZ-X710 microscope. Organoid growth and formation efficiency were assessed for each condition at endpoint (Fig. 1c, Extended Data Fig. 1k).

#### Histology and immunofluorescence detection of proteins

Intestinal tissues were fixed in 4% paraformaldehyde at 4 °C overnight and washed with PBS to remove excess paraformaldehyde. Tissues were dehydrated by passing through an ethanol gradient, embedded in paraffin, and sectioning was done at 5 μm thickness. Sections were deparaffinized, rehydrated, and stained with the alkaline phosphatase substrate nitro blue tetrazolium/5-bromo-4-chloro-3-indolyl-phosphate (NBT/BCIP; Sigma, B5655) or with alcian blue (Mucin). For immunofluorescence, 10 mM sodium citrate buffer (pH 6.0) was used for antigen retrieval and sections were incubated overnight with following primary antibodies at 4 °C in PBS with 0.1% BSA: anti-Ki67 (Thermo Fisher, 14-5698-80), anti-GFP (Santa Cruz Biotechnology, sc-9996), anti-H3K27me3 (Diagenode, C15410195), anti-RFP (Rockland, 600-401-379), and anti–β-catenin (Cell Signaling Technology, 8480S). The following day, sections were incubated with Alexa Fluor–conjugated secondary antibodies (anti-rabbit, anti-mouse, or anti-donkey IgG; Alexa Fluor 488, 555, or 647; Thermo Fisher; 1:1000 dilution). DAPI was used for nuclear labeling pripr to imaging on fluorescence microscope.

#### Whole-intestine endogenous fluorescence imaging

Intestinal tissues from the duodenum to the cecum were collected, fixed overnight in 4% paraformaldehyde at 4 °C, and washed five times with PBS. Tissues were then incubated in 30% sucrose in PBS overnight at 4 °C for cryoprotection. Samples were embedded in optimal cutting temperature (OCT) compound and stored at −80 °C until sectioning. Cryosectioning was performed using a Leica CM3050 S cryostat to obtain 7 μm tissue sections. Sections were mounted using VECTASHIELD Antifade Mounting Medium with DAPI (Vector Laboratories, H-1200-10) to visualize nuclei. Endogenous fluorescence was used to identify *Lgr5^+^* and *Lgr5^‒^* cell populations, detected via FITC and TRITC channels, respectively. Whole-intestine imaging was performed using a Zeiss Axioscan.Z1 slide scanner (Extended Data Fig. 1b).

#### hybrid organoid based clonal-competition assays

*Lgr5^+^* cells were isolated by FACS from day 7 *Apc^‒/‒^* and *Apc^‒/‒^ Eed^‒/‒^* intestinal tissues and co-cultured at a density of 100,000 cells per well in 24-well ultra-low attachment aggregate plates (STEMCELL Technologies) to allow spheroid formation. After 24 hours, heterogeneous spheroids were embedded in 30 μl Matrigel (Corning, 356231) and cultured in mutant intestinal organoid medium (standard intestinal medium lacking R-spondin-conditioned medium and Y-27632). Organoids were maintained in culture for 3 days and imaged every day using a Keyence BZ-X710 microscope. Quantification was performed on all days by measuring changes in the circumference of each organoid fraction relative to their starting size (Fig. 7g, Extended Data Fig. 7i).

#### Native-ChIP-seq libraries

Native-ChIP-seq was performed as previously described^20, 50^. In brief, cells were lysed in 0.1% Triton X-100 and 0.1% Sodium Deoxycholate with protease inhibitor cocktail. Chromatin in the cell lysate was digested using Microccocal nuclease (MNase, New England BioLabs, M0247S) at room temperature for 5 min and 5 ul 0.25 mM EDTA was added to stop the reaction. The digested chromatin were incubated with anti-IgA magnetic beads (Dynabeads,Themo Fisher, 10001D) for 2 h for pre-clearing and then incubated overnight with antibody-bead complexes with 0.5ug of antibodies against H3K27me3 and H3K27ac (Diagenode, C15410195 and C15410046, respectively) in immunoprecipitation (IP) buffer (20 mM Tris-HCl pH 7.5, 2 mM EDTA, 150 mM NaCl, 0.1% Triton X-100, 0.1 % Sodium Deoxycholate) at 4 °C. IPs were washed 2 times by Low Salt (20 mM Tris-HCl pH 8.0, 2 mM EDTA, 150 mM NaCl, 1% Triton X-100, 0.1% SDS) and High Salt (20 mM Tris-HCl pH 8.0, 2 mM EDTA, 500 mM NaCl, 1% Triton X-100, 0.1% SDS) wash buffers. IPs were eluted in elution buffer (1% SDS, 100 mM Sodium Bicarbonate) for 1.5 h at 65 °C. Histones were digested by Protease (Qiagen 19155) for 30 min at 50 °C and DNA fragments were purified using Sera Mag magnetic beads in 30% PEG. Illumina sequencing libraries were generated as previously described by end repair, 3′ A-addition, and Illumina sequencing adaptor ligation (New England BioLabs, E6000B-10). Libraries were then indexed and PCR amplified (8 cycles) and sequenced on Illumina NovaSeq 6000 sequencing platform at Genewiz (Azenta Life Science).

#### Whole genome bisulfite sequencing (WGBS) libraries

Genomic DNA was extracted using the DNeasy Blood & Tissue Kit (Qiagen, 69504) according to the manufacturer’s instructions. WGBS libraries were prepared from 25 ng of purified DNA using the Pico Methyl-Seq Library Prep Kit (Zymo Research, D5455), following the manufacturer’s protocol with the modification of reducing the number of PCR amplification cycles to a total of 8. Libraries were sequenced on an Illumina HiSeq 2500 platform (Novogene) to generate paired-end reads.

#### CUT&RUN

CUT&RUN was performed using the ChIC/CUT&RUN Assay Kit (Active Motif, 53180) following the manufacturer’s protocol with minor modifications. Freshly isolated epithelial cells were resuspended in permeabilization buffer (ChIC/CUT&RUN Wash Buffer supplemented with 0.01% digitonin and 1× protease inhibitor cocktail) and incubated on ice for 5 minutes. Cells were then bound to Concanavalin A–coated magnetic beads at room temperature for 10 minutes. Bead-bound cells were resuspended in 180 μl of antibody buffer (permeabilization buffer with 2 mM EDTA) containing 0.5 μg of anti-Cdx2 (Cell Signaling, D11D10) and incubated overnight at 4 °C. Nuclear translocation of micrococcal nuclease (MNase) was performed at 4 °C for 1 hour, followed by chromatin digestion at 4 °C for 30 minutes on a nutator. Fragment release was carried out at 4 °C for 30 minutes, and DNA was purified using SeraMag magnetic beads in 30% PEG-8000. Libraries were prepared as described in the Native ChIP–seq section.

#### Single-cell multiome libraries (combined scRNA and ATAC)

Libraries were prepared using the Chromium Next GEM Single Cell Multiome ATAC + Gene Expression kit (10x Genomics) according to the manufacturer’s protocol. Briefly, nuclei were isolated and transposed using Tn5 transposase, followed by reverse transcription and barcoding within individual Gel Beads-in-Emulsion (GEMs). Libraries for both ATAC and gene expression were amplified and purified separately. Final libraries were quantified using Qubit dsDNA HS Assay (Thermo Fisher Scientific) and quality assessed with an Agilent Bioanalyzer. Sequencing was performed on an **Illumina** NextSeq 6000 using paired-end reads with a read configuration of 28 bp (Read 1), 90 bp (Read 2), and 10 bp (i7 index).

### QUANTIFICATION AND STATISTICAL ANALYSIS

#### RNAseq

Raw RNA-seq reads were aligned to the mouse genome (GRCm38, GENCODEv25) using STAR aligner v2.7.8a^50, 51^ in two-passed mode followed by assessment to determine per-base sequence quality, per-read GC content (∼50%), comparable read alignments to +/- strands, exon vs intron read distributions, and 3’ bias. Transcript levels were expressed as read counts using HTSeq v0.13.5 and normalized across libraries using DESeq2,^52^ followed by conversion into reads per kb of transcript length per 1M mapped reads (RPKM). We determined differential expression between samples using DESeq2, with false-discovery rate (FDR) as indicated in the text. All alignments and initial processing steps were run using the Snakemake workflow management system (v6.6.1). Cartoon Illustrations were created with BioRender.

#### ChIP-seq

ChIP-seq data were trimmed using trim-galore (v0.6.6) to remove adapters and low-quality reads. Trimmed sequences were aligned using bwa-mem (v0.7.17)^53^ with default parameters. Non-uniquely mapped reads, duplicate reads, and reads with a MAPQ score of less than 5 were removed using Samtools (v1.17).^54^ Peak calling was done using MACS (v2.2.7.1)^55^ with a q-value cutoff of 0.01 in sharp mode for H3K27ac and broad mode for H3K27me3. BigWig files were generated using deeptools (v3.5.1)^56^ with a bin size of 10, smooth length of 30, and quantile normalized using HayStack. Bedtools (v2.30.0)^57^ was used to removed blacklisted regions defined by ENCODE blacklist.

#### Analysis of DNA methylation data

Whole genome bisulfite data were aligned to mm10 in non-strand specific mode using Bismark v0.23.0. Coverage from both strands for each CpG were combined and fractional methylation was calculated for each CpG using custom AWK command. Differential single CpGs were identified using fisher exact test (p < 0.01). Differential CpG with at least 20% difference in fractional methylations and within 300bp of one another were merge using custom AWK command. Differential methylated regions (DMR) with less than 40% methylation were discarded.

#### Motif enrichment analysis

Motif enrichment was conducted using HOMER^41^ findMotifGenome with default parameters (Fig 4h).

#### Single-cell Multiom data analysis

scMultiome data were aligned and processed using Cell Ranger Arc (10x Genomics, version 2.0.0). Downstream analyses were performed using Seurat v4 and Signac for stringent quality control and data integration of single-cell RNA-seq (scRNA-seq) and single-nucleus ATAC-seq (snATAC-seq) datasets. For the scRNA-seq data, cells were retained if they met the following criteria: >12,000 unique molecular identifiers (UMIs) per cell, >1,000 detected genes per cell, mitochondrial ratio <0.35, and expression novelty >0.8. For the snATAC-seq data, nuclei were retained if they met the following criteria: percentage of reads in peaks >40%, ATAC fragments per cell >1,000, blacklist fraction <0.05, TSS enrichment >2, and nucleosome signal <4. After filtering, a total of 8,419 high-quality cells were retained for downstream analysis — including 4,511 cells from wild-type (WT) and 3,908 cells from day 14 *Apc⁻/⁻* samples. The Seurat package was used for data normalization, dimensionality reduction (PCA and UMAP), clustering, and visualization of cell states and identities.

#### Pseudotime Trajectory Analysis

Pseudotime analysis was performed using Monocle 2 to infer transcriptional trajectories and lineage relationships. The normalized expression matrix from Seurat was imported into Monocle, and dimensionality reduction was carried out using UMAP after principal component analysis (PCA). Cells were ordered along trajectories using “learn_graph” and “order_cells” functions based on stem-enhancer associated genes. Branch points were identified to delineate transcriptional transitions between stem and non-stem cells. The pseudotime ordering was further validated by the expression of canonical stem markers (Fig. 3c-e, Extended Data Fig. 3g).

#### Calculation of enhancer activity score representing the balance of activating (H3K27ac) and repressive (H3K27me3) marks

Peaks for H3K27me3 and H3K27ac were called using MACS2 with default parameters. Genomic regions of 2000 bp with peaks of both or either of the marks were identified. Regions with peaks of both were centered on the summit of the H3K27ac peaks and the rest on the peak summits for the individual mark. Regions with fewer than 30 reads for both the marks were removed as low signal loci. For each region, activation (H3K27ac) and repression (H3K27me3) signals were quantile-normalized to each other. A weighted signal score, which we refer to as, “action score”, was calculated as the ratio of H3K27ac signal to the combined signal of both marks (Fig. 1e, Extended Data Fig. 2a-e).

Enhancer activity score (AS) = H3K27ac_normalized count / (H3K27me3_ normalized count + H3K27ac_ normalized count), which ranged from 0 (most inactive and repressed enhancers, without any H3K27ac) to 1 (most active enhancers, without any H3K27me3). Significant changes in the enhancer activity scores were calculated using Fisher’s exact test and enhancers with a *P* < 0.0001 and an absolute change of 0.3 were considered to have changed the activity significantly. Regions with less than 0.3 change in activation were mostly below p value threshold. Replicates of H3K27me3 and H3K27ac data showed Pearson correlation of > 0.7 in both *WT* and *Apc^‒/‒^* cells ensuring applicability of the Fisher exact test.

|  | H3K27ac | H3K27me3 | SUM |
| --- | --- | --- | --- |
| WT | a | b | a + b |
| APC | c | d | c + d |
| SUM | a + c | b + d | a + b + c + d |

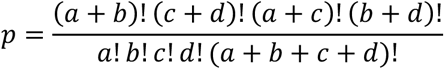

## Data availability

The sequencing data reported in this study is accessible through Gene Expression Omnibus (GEO). The accession number for the data generated in this study are RNAseq: GSE31795, ChIP-seq: GSE319207 and Cut & Run: GSE317495. The multiomics data is accessible through GSE315065.

## Acknowledgments

This work was supported by NIH grants K01DK113067, R03DK134799, Donald E. and Delia B. Baxter Foundation Fellowship Award, Tower Cancer Research Foundation Fellowship Award, and The Uri and Karen Geiger Cancer Metastasis Research and Innovation Fund to U.J. A.L was supported by Canadian Institute of Health Research (CIHR) Post-Doctoral Fellowship and California Institute of regenerative Medicine (CIRM) Post-Doctoral Fellowship. We would like to thank Dr. Kwok-Kin Wong (NYU Langone Health) for providing the Col1a-LSL-Ezh2 mice.

## Contributions

A.L. U.J. and R.A.S conceptualized the study design. A.L. and G.Y. performed the analysis. A.L., S.S., S.R. and Y.J.K. conducted experimental work. A.L., U.J. wrote the manuscript, A.L., U.J, and R.A.S. edited the manuscript. All authors read and approved the final manuscript.

---

**Extended Data Figure 1.**
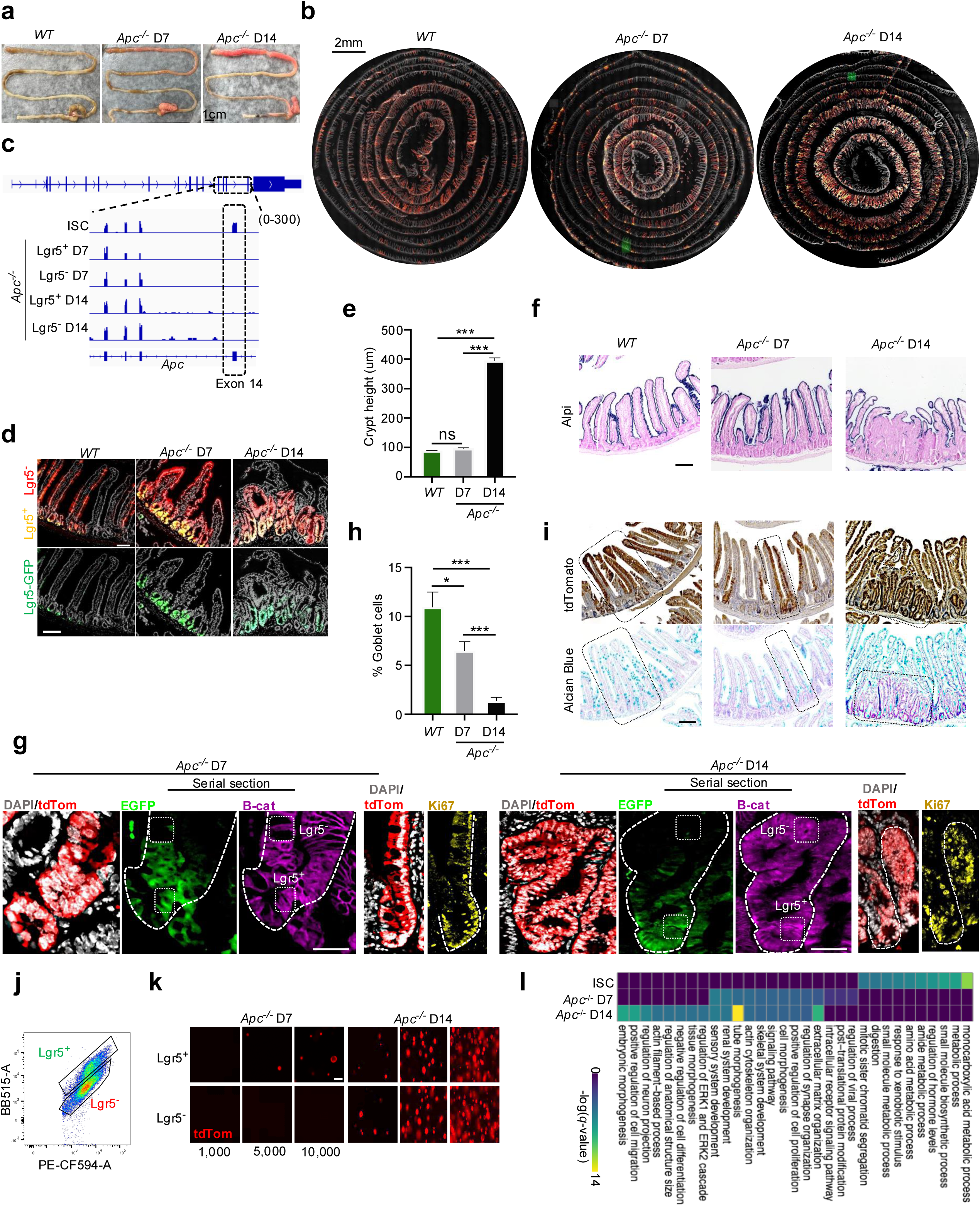
**a.** Gross intestinal morphology from *WT*, *Apc^‒/‒^* D7, and *Apc^‒/‒^* D14 mice showing progressive polyp growth. Scale bar, 1 cm. **b.** Whole-mount fluorescence imaging showing expansion of transformed crypt domains in *Apc^‒/‒^* tissues over time (Lgr5^+^: GFP^+^/tdTomato^+^, yellow; Lgr5^‒^: tdTomato^+^, red). Scale bar, 2 mm. **c.** Browser tracks of RNA-seq confirming deletion of Apc exon 14 in *Apc^‒/‒^* D7 and D14 Lgr5^+^ and Lgr5^‒^ cells. **d.** Fluorescence imaging of crypt–villus units displaying expansion of Lgr5^+^ cells (green/yellow) and emergence of transformed Lgr5^‒^ cells (red) after Apc loss. Scale bar = 100um. **e.** Quantification of crypt height in *WT* and *Apc^‒/‒^* mice (n > 3 mice; mean ± s.e.m.; ***p < 0.001; ns, not significant). **f.** Alkaline phosphatase (Alpi) staining showing altered epithelial architecture and loss of enterocyte differentiation in *Apc^‒/‒^*intestines at D14. Scale bar = 100um. **g.** Serial-section immunofluorescence for tdTomato, EGFP, β-catenin, and Ki67, showing minimal nuclear β-catenin and MKi67 (proliferation) signal in Lgr5^‒^ cells at D7, followed by their gain into the non-Lgr5 cells by D14. Scale bar = 50um. **h.** Quantification of goblet cell frequency across conditions (mean ± s.e.m.; n = 3 mice; \**p* < 0.05, \*\*\**p* < 0.001) **i.** Immunohistochemistry for tdTomato (Cre-recombined cells) and Alcian blue staining for goblet cells (serial sections) at D7 and D14, showing loss of differentiation at D14. Scale bar = 100um. **j.** Representative flow cytometry gating for FACS isolation of Lgr5^+^ and Lgr5^‒^ epithelial cells from *WT* and *Apc^‒/‒^* mice. **k**. Fluorescent micrographs of organoids formed from FACS-isolated Lgr5^+^ and Lgr5^‒^ *Apc*^‒/‒^ cells (D7 and D14) seeded at 1,000, 5,000, or 10,000 cells, showing endogenous tdTomato expression at day 6. Scale bar = 50um. **l.** gene ontology analysis of differential expressed genes across ISCs and *Apc^‒/‒^* Lgr5^+^ cells at D7 and D14.

**Extended Data Figure 2.**
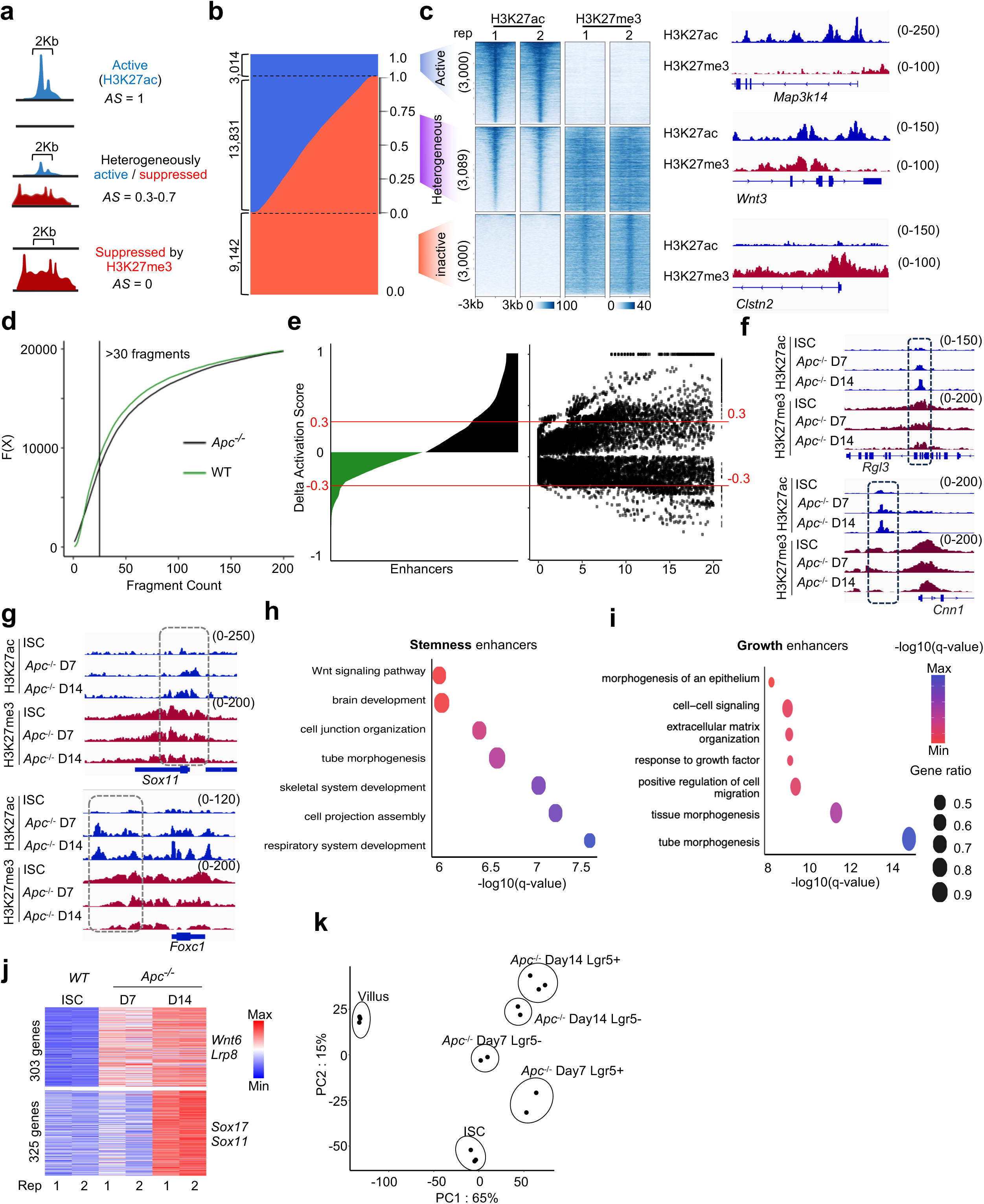
**a.** Schematic of enhancer-state classification based on activity score (AS) derived from H3K27ac and H3K27me3 signal (over 2 kb window): active (AS = 1), heterogeneously active (AS = 0.3–0.7), and repressed (AS = 0). **b.** Distribution of enhancers across AS categories in ISCs. **c.** Heatmaps of H3K27me3 and H3K27ac signals in ISC at 3,089 plastic, and 3,000 randomly selected active and repressed enhancers, with representative browser tracks (*Map3k14*, *Wnt3*, and *Clstn2*) shown at right illustrating enhancer categories. **d.** Cumulative H3K27ac fragment distributions at MACS2-called enhancers marked by H3K27ac or H3K27me3 in ISCs and *Apc^‒/‒^* Lgr5^+^ cells at day 14; vertical line marks fragment cutoff used for selecting enhancers for downstream analysis. **e.** Histogram of ΔAS between ISCs (green) and *Apc^‒/‒^* day 14 (black), and volcano plot identifying significantly activated (ΔAS > 0.3) and repressed (ΔAS < –0.3) enhancers. **f.** Genome browser views showing activation of stemness- enhancers (of genes *Rgl3* and *Cnn1*) in *Apc^‒/‒^* intestines. **g.** Genome browser views showing activation of growth- enhancers (of genes *Sox11* and *Foxc1*) in *Apc^‒/‒^* intestines. **h,i.** Gene-set enrichment analysis of genes linked to stemness (h) enhancers highlights Wnt signaling and developmental morphogenesis pathways, while growth (i) enhancers are linked to cell migration and proliferation related processes. **j.** Heatmap showing expression (log2RPKM+1) of genes linked to (gene TSS within 25kb) stemness or growth enhancers; expression values from 2 replicates of ISCs and *Apc^‒/‒^* Lgr5^+^ cells at D7 and D14 are represented. **k.** Principal component analysis of gene expression (mRNA-seq) data from ISCs, villus cells, and *Apc^‒/‒^* Lgr5^+^/Lgr5^‒^populations from D7 and D14.

**Extended Data Figure 3.**
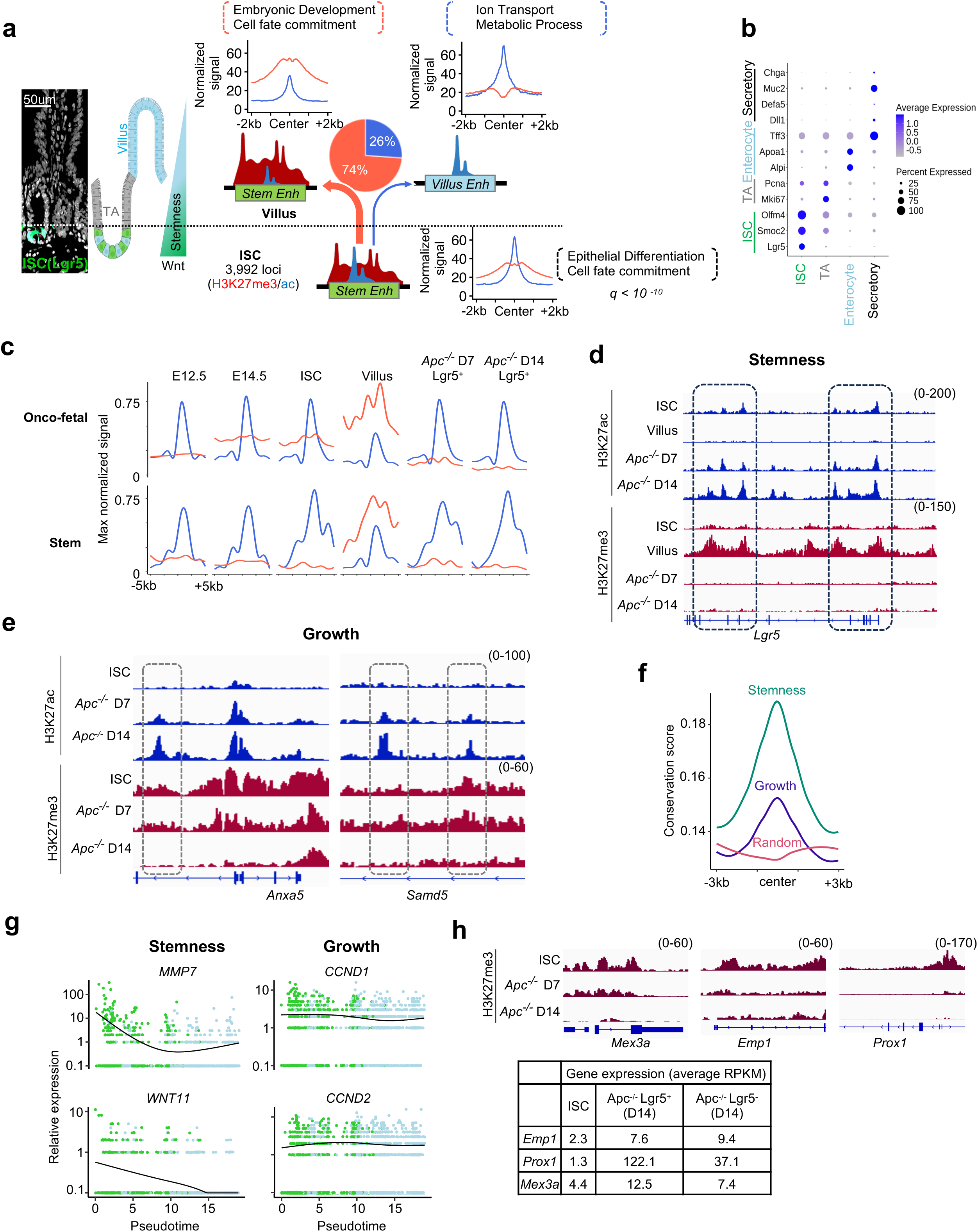
**a.** Fluorescence micrograph of the mouse intestinal crypt–villus axis showing endogenous EGFP signal from Lgr5^+^ ISCs, with schematic of ISC, transit-amplifying (TA), and villus compartments along the Wnt and corresponding stemness gradient. Schematic of 3,992 ISC enhancers co-marked by H3K27ac and H3K27me3, most of which (74%) lose activity and gain repression upon villus differentiation, while a minority (26%) gain activity. Profile plots show cumulative H3K27ac (blue) and H3K27me3 (red) signals in ISCs and villus epithelium for these enhancers. Gene Ontology enrichment of ISC enhancers either repressed or activated in villus cells (q < *10⁻¹⁰*). **b.** Dot plots showing expression levels and percent of cells expressing lineage marker genes used to define clusters in the single-cell multiome data. **c.** Profile plots of H3K27ac and H3K27me3 signals (normalized to max) at onco-fetal and stem gene promoters^26^ across fetal (E12.5, E14.5), adult ISCs and villus cells, and *Apc*^‒/‒^ Lgr5^+^ cells at D7 and D14, showing reactivation of fetal H3K27me3-protected programs during early transformation. **d.** Genome browser tracks showing enhancer hyperactivation at the *Lgr5* locus in *Apc*^‒/‒^ Lgr5^+^ cells at day 7 and day 14 compared to repression in differentiated villus cells in *WT* intestine. **e.** Genome browser tracks showing increased H3K27ac and loss of H3K27me3 at *Anxa5* and *Samd5* enhancers in *Apc*^‒/‒^ samples relative to ISCs. **f.** Sequence conservation profiles (±3 kb from enhancer center) showing that stemness enhancers are the most evolutionarily conserved across vertebrates relative to growth or 3,000 random villus-specific enhancers. **g.** Expression of stemness (*MMP7*, *WNT11*) and growth (*CCND1*, *CCND2*) enhancer-linked genes plotted along pseudotime in stem-like (green) and non-stem (blue) tumour cells. **h.** Genome browser tracks showing loss of H3K27me3 at *Emp1, Prox1,* and *Mex3a* gene enhancers in *Apc*^‒/‒^ samples along with a table showing increased expression of these genes: average RPKM values of mRNA-seq data are shown for Lgr5^+^ and Lgr5^‒^ cells.

**Extended Data Figure 4.**
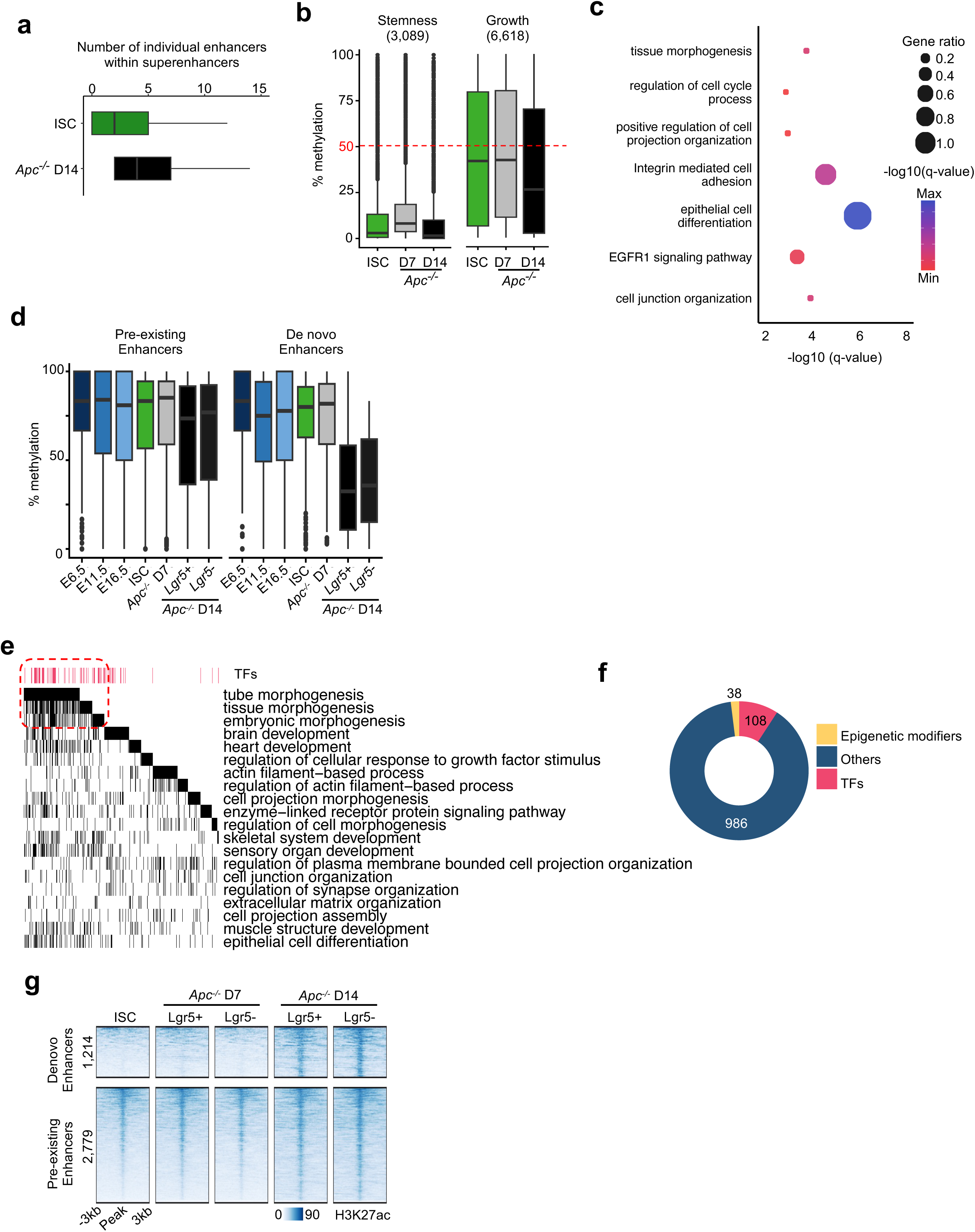
**a.** Bar plot showing the average number of active enhancers per superenhancer in ISCs and *Apc^‒/‒^* D14 cells. **b.** Box and whisker plot showing hypomethylation at stemness and growth enhances with little change during cell transformation in *Apc^‒/‒^* D7 and D14 cells. **c.** Gene ontology analysis of genes associated with hypomethylated *de novo* enhancers highlighting enrichment in pathways related to tissue morphogenesis and epithelial organization. **d.** Box and whisker plot showing high methylation levels at *de novo* and pre-existing enhancers during development and in adult intestinal epithelial cells. *De novo* enhancers show substantial hypomethylation in *Apc^‒/‒^*D14 cells. **e.** Binary heatmap of genes identified in the gene ontology analysis that lose H3K27me3 at their promoters and are associated with stemness enhancers, showing enrichment of transcription factors linked to development and morphogenesis. **f.** Donut chart summarizing number of TF, epigenetic modifier, and other genes linked to with stemness enhancers and significant loss of H3K27me3 at their promoters. **g.** Heatmaps of H3K27ac signals at *de novo* and pre-existing enhancers formed within superenhancers in *Apc^‒/‒^* D14 across ISC, Lgr5^+^ and Lgr5*^‒^* at D7 and D14.

**Extended Data Figure 5.**
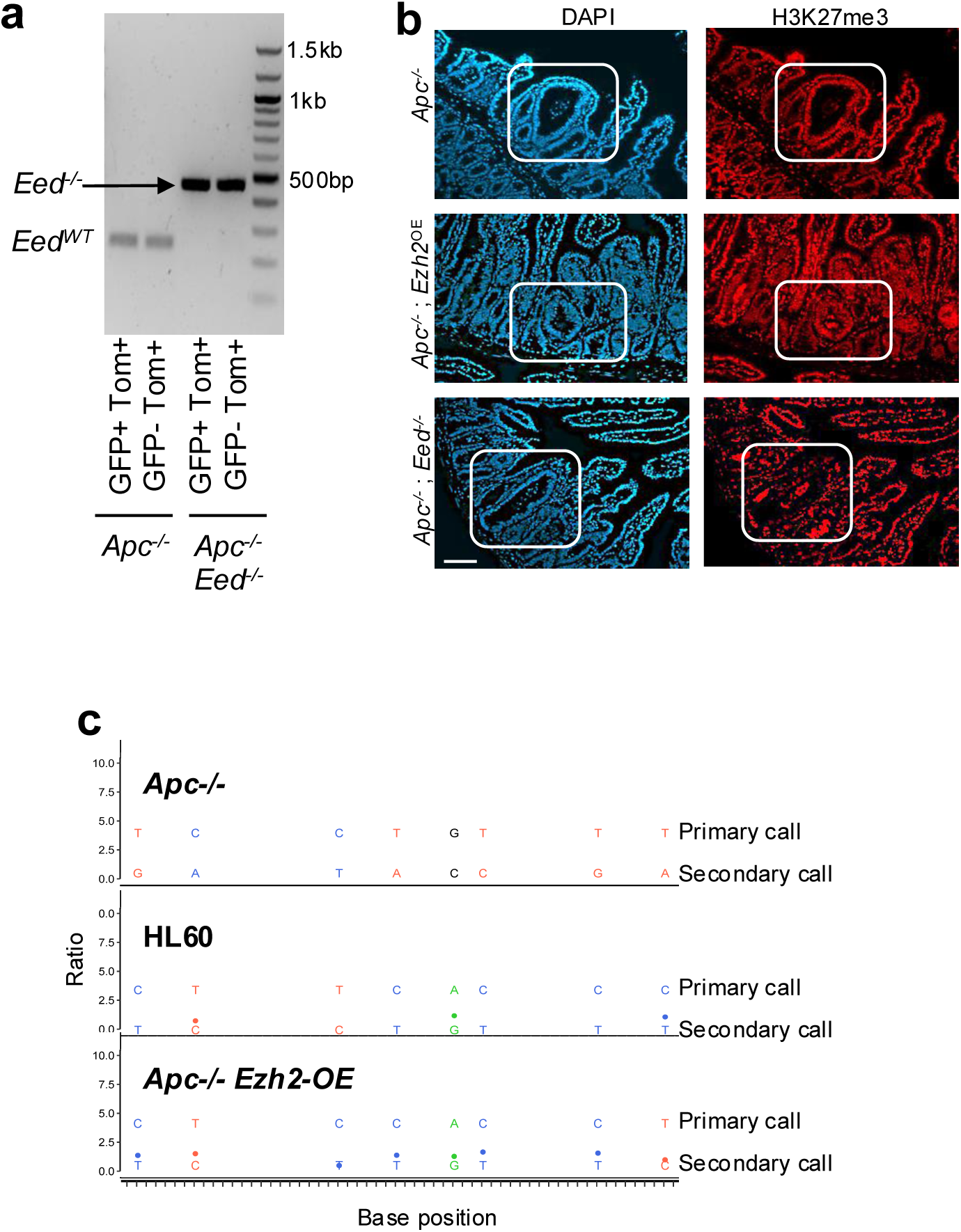
**a.** PCR of genomic DNA from FACS isolated Lgr5^+^ and Lgr5^‒^ cells from *Apc^‒/‒^ Eed^‒/‒^* epithelium confirms *Eed* deletion by generating a larger amplification product. **b.** Immunofluorescence staining for DAPI (blue) and H3K27me3 (red) in intestine of *Apc^‒/‒^*, *Apc^‒/‒^Eed^‒/‒^* and *Apc^‒/‒^ Ezh2*^OE^ mice. Boxed regions highlight representative crypts showing reduction in H3K27me3 upon loss of *Eed* and gain upon overexpression of *EZH2*. Scale bar = 100um. **c.** Sanger sequencing of EZH2 mRNA showing matching bases between human mRNA (from HL60 cells) and Lgr5^+^ cells from *Apc^‒/‒^ Ezh2*^OE^ cells; *Apc^‒/‒^* cells do not show sequence similarity to human mRNA.

**Extended Data Figure 6.**
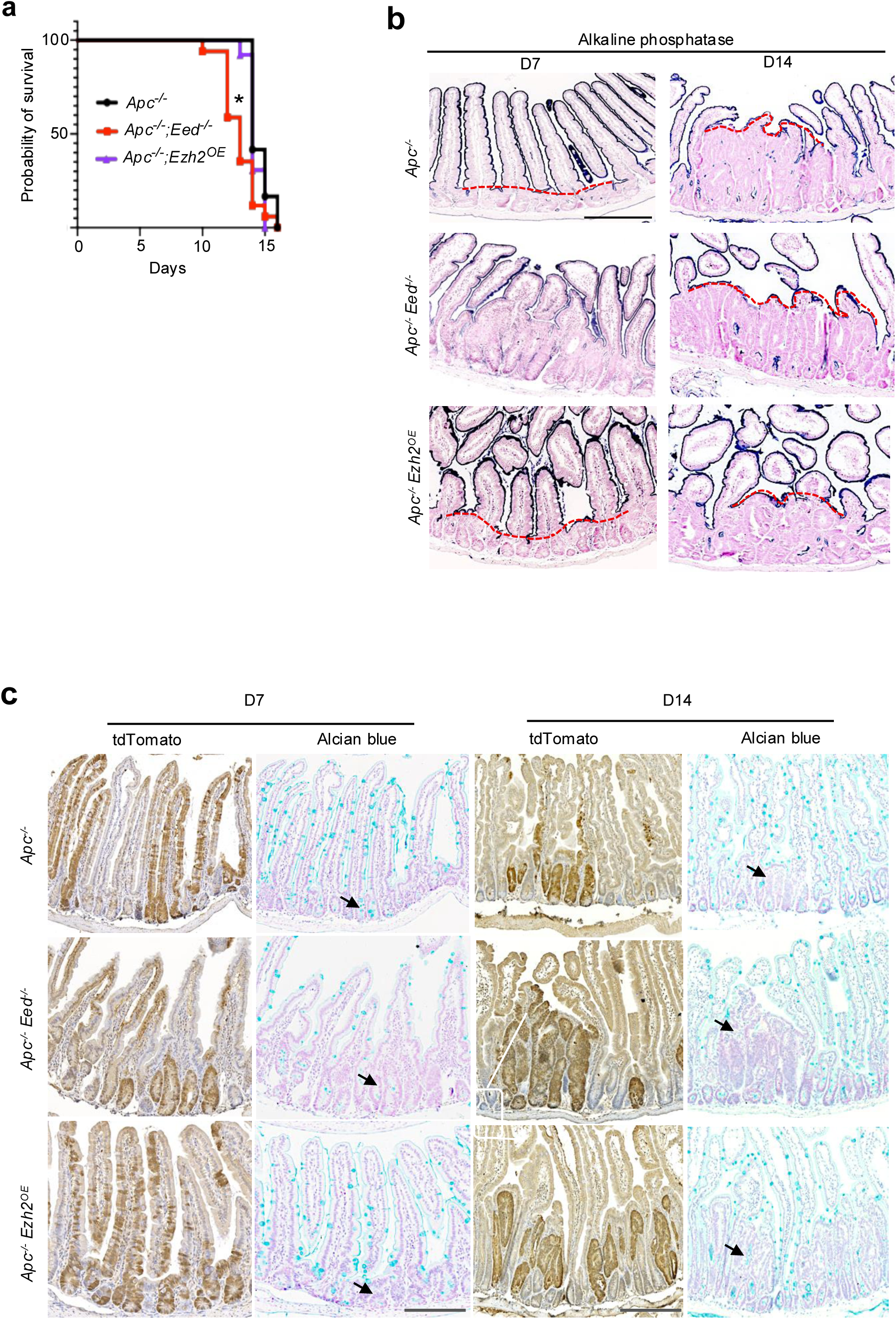
**a.** Kaplan–Meier survival curves for *Apc^‒/‒^*, *Apc^‒/‒^ Eed^−/–^*, and *Apc^‒/‒^Ezh2^OE^*mice showing reduced survival in *Apc^‒/‒^ Eed^−/–^* mice (n > 10, *p* < 0.05). **b.** Alkaline phosphatase staining (enterocytes on villus) at D7 and D14 showing altered differentiation zone boundaries (dashed red lines). *Apc^‒/‒^ Eed^‒/‒^*mice show accelerated expansion of stem cells, pushing the differentiated cell zone farther from the crypt bottom. Scale bar = 200um. **c.** Immunohistochemistry for tdTomato (all transformed cells) and goblet cells (alcian blue staining of Mucin) at D7 and D14. *Apc^‒/‒^ Eed^‒/‒^* adenomas show earlier reduction in goblet cell differentiation at D7 compared with PRC2-intact tumours (arrows). Scale bar = 200um.

**Extended Data Figure 7.**
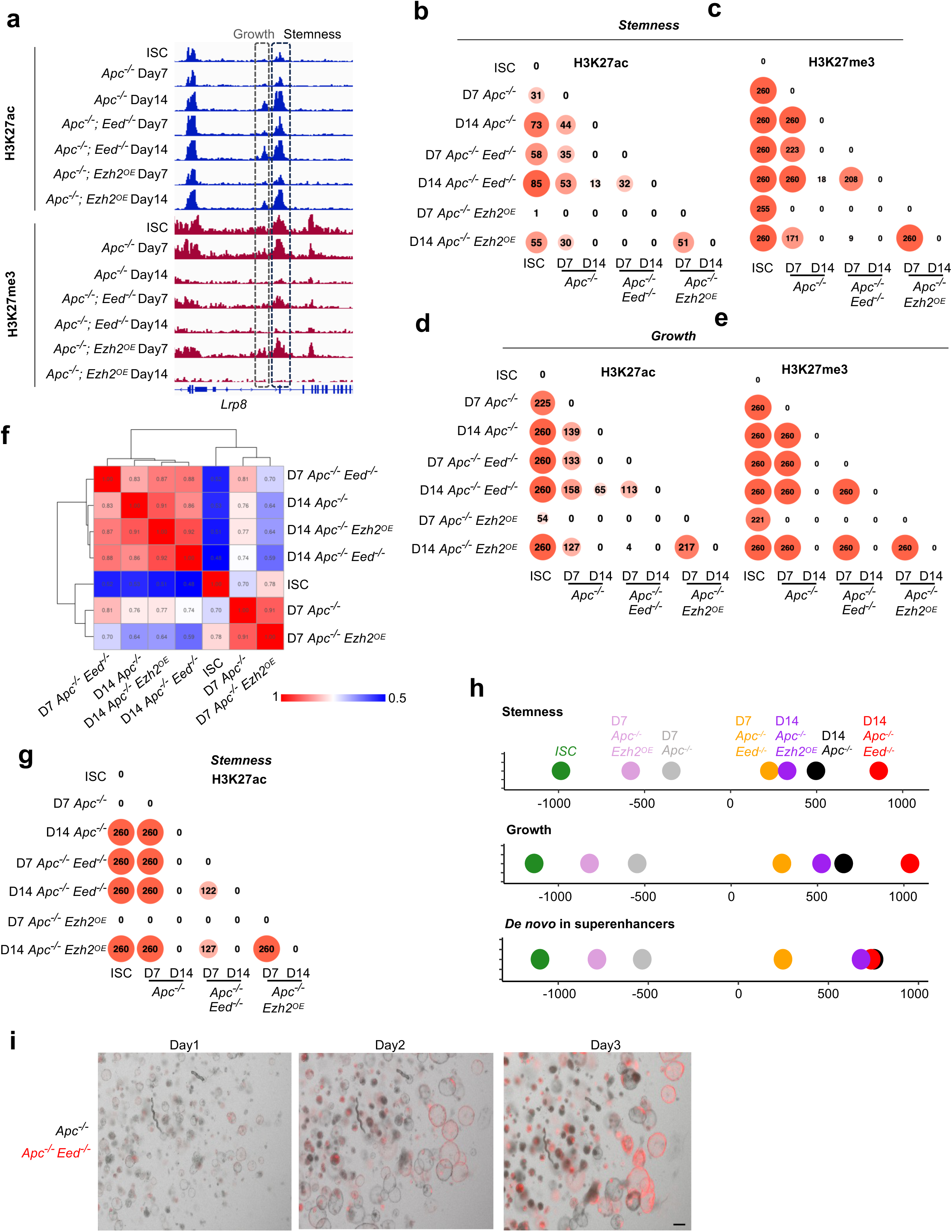
**a.** Genome browser tracks showing H3K27ac and H3K27me3 signals in Lgr5^‒^ cell populations across indicated samples, highlighting activation of growth-associated enhancers in *Apc^‒/‒^*; *Eed^‒/‒^* D7. **b-e.** Dot plots representing significance of differences in H3K27ac **(b, d)** or H3K27me3 **(c, e)** signal across each tumour sample relative to ISCs (one-sided Kolmogorov-Smirnov *p*-value) at stemness and growth enhancers. **f.** Heatmap showing correlation of H3K27ac signal across indicated samples at enhancers located within *Apc^‒/‒^* D14 superenhancers. **g.** Dot plots representing significance of differences in H3K27ac signal across each tumour sample relative to ISCs (one-sided Kolmogorov-Smirnov *p*-value) at *de novo* enhancers within *Apc^‒/‒^* superenhancers. **h.** Uniscale reduction analysis illustrating relative similarity of Lgr5^+^ cell states based on stemness and growth enhancer activity across all mouse models. *Apc^‒/‒^ Eed^‒/‒^* cells show faster activation of both type of enhancers as well as higher level of activation while overexpression of *Ezh2* (*Apc^‒/‒^ Ezh2*^OE^) leads to slower and reduced stemness gain and growth. **i.** Overlay of brightfield and fluorescence microscopic images showing hybrid organoids (*Apc^‒/‒^*, uncolored; *Apc^‒/‒^ Eed^‒/‒^*, red) at days 1, 2, and 3, illustrating accelerated growth upon PRC2 loss. Scale bar = 50um.

